# Cryo- EM structure of ribosome from pathogenic protozoa *Entamoeba histolytica,* reveals unique features of its architecture

**DOI:** 10.1101/2024.04.29.591517

**Authors:** Shivani Sharma, Shalini Mishra, Samudrala Gourinath, Prem S. Kaushal

## Abstract

*Entamoeba histolytica*, an anaerobic parasite protozoan, is the causative agent of amoebiasis, the bloody diarrhea, and liver abscesses in humans. Amoebiasis is more predominant in tropical areas with poor sanitation conditions, and it remains the fourth leading cause of death due to a protozoan infection. *E. histolytica* life cycle spans between an infective ‘cyst stage’ and an active disease-causing ‘trophozoite stage’. We have determined cryo-EM structures of *E. histolytica* ribosomes, large subunit (LSU), 53S ribosome at 2.8 Å resolution and associated, 75S ribosome at 3.3 Å resolution, isolated from its trophozoite stage. The overall core of the ribosome is conserved. However, the periphery has evolved with *Entamoeba* specific unique features. The most notable features are, the presence of the rRNA triple helix near the peptide exit tunnel on LSU and the co-evolution of rRNA expansion segments and extensions in r-proteins. To the best of our knowledge, this structure reports the presence of RNA triple helix in the ribosome for the first time.

## Introduction

*Entamoeba histolytica*, as its name suggests (histo-lytic = tissue destroyer), is an anaerobic parasite responsible for an intestinal infection that results in blood diarrhea known as invasive amoebiasis or invasive colitis (Stanley 2003). Occasionally, the parasite reaches into the bloodstream and spreads through the body, most frequently ending up in the liver, where it causes liver abscesses. In rare cases, it reaches the lung, brain, and spleen and causes abscesses, which can be fatal if untreated (Nespola et al., 2015; Ryan *et al*., 2004; Stanley, 2003). *E. histolytica* infects approximately 35-50 million people worldwide and roughly kills 55,000 people each year (https://www.cdc.gov/dpdx/amebiasis/index.html ) (Shirley et al., 2018). The life cycle of *E. histolytica* spans, the active disease-causing stage, ‘trophozoites’, which exist only in the host, and the inactive stage ‘cysts’, which survive outside the host. Infection occurs via ingestion of mature cysts from fecally contaminated food, water, or hands (Shirley et al., 2018).

Earlier studies showed that ribosomes from *Entamoeba invadens*, a non-pathogenic model organism for *E. histolytica*, assemble in a rod-shaped helical structure known as chromatoid bodies (Barker & Deutsch, 1958; Morgan & Uzman, 1966; Morgan, 1968). Meanwhile, for the first time, the protein synthesis in *Entamoeba* trophozoites was first reported by Carter et al., 1967. The purified ribosomes form helices of the cyst, and polysomal bound from growing trophozoites appeared identical (Kusamrarn et al., 1975). However, the crude cyst extract was inefficient in protein synthesis, and it became efficient only after adding the soluble fractions from trophozoites (Kusamrarn et al., 1975). Emetine was introduced as an amoebicidal drug that binds to the *Entamoeba histolytica* ribosome and inhibits protein synthesis (Entner & Grollmann, 1972; Entner, 1979). However, within two decades, *E. histolytica* mutants that were resistant to emetine were also reported (Orozco et al., 1985). Currently, ribosome targeting aminoglycoside antibiotic ‘paromomycin’ along with drugs are used as amoebicidal.

*Entamoeba* possesses reduced ribosomes and its monosome, large subunit (LSU), and small subunit (SSU) sediments at 75S, 53S, and 36S, respectively (Price et al., 1983). It was reported that *Entamoeba* ribosome possesses five species of rRNA: 25S, 17S, 5.8S, 5S, and 4S (Albach et al., 1984). Most eukaryotic ribosomes, monosome, LSU, and SSU sediments, at the 80S, 60S, and 40S, respectively, and possess four rRNA species 28S, 18S, 5.8S and 5S. The high-resolution ribosome structures from other protozoans: *Trypanosoma brucei* (Hashem et al., 2013), *Plasmodium falciparum* (Wong et al., 2014), *Leishmania donovani* (Shalev-Benami et al., 2017), *Trichomonas vaginalis* and *Toxoplasma gondii* (Li et al., 2017), *Euglena gracilis* (Matzov et al., 2020) *Giardia lamblia* (Hiregange et al., 2022) (Majumdar et al., 2023); Eiler et al. 2024) have revealed the species-specific unique features of their architecture and protein synthesis. The atomic details for *E. histolytica* ribosome architecture are absent as no structure is available.

Here, we report the cryo-EM single particle reconstruction structure of *E. histolytica* 53S, ribosome large subunit, at 2.8Å resolution and 75S ribosome at 3.3 Å resolution. The structures provide several unique features specific to *E. histolytica* ribosome architecture, such as the rRNA triple helix structure, *Entamoeba* specific expansion segments in rRNA, and unique insertion in r-proteins. With this structure, we also corrected and reannotated rRNA length and r-proteins.

## Results and Discussion

### E. histolytica ribosome

*The E. histolytica* ribosomes purified through sucrose density gradient centrifugation (SDGC) showed three prominent peaks (Supplementary Fig. S1a). The fractions analyzed on 1% agarose gel showed two prominent bands after ethidium bromide staining, suggesting that it contains ribosomal RNA (rRNA), most likely the 25S and 17S rRNAs (Supplementary Fig. S1a). *E. histolytica* ribosome consists of 25S, 17S, 5.8S, 5S, and 4S rRNA (Albach et al., 1984), which are slightly condensed as compared to human rRNAs; 28S, 18S, 5.8S, and 5S (Natchiar et al., 2017). The fractions corresponding to the peaks P1, P2, and P3(Supplementary Fig. S1a), analyzed through negative staining electron microscopy, clearly showed that peak P1 contains irregular shape and size particles (Supplementary Fig. S1b), which might be degraded ribosomes. Peak 2 contains large ribosomal subunit (LSU), which is 53S in *Entamoeba* (Price et al., 1983) (Supplementary Fig. S1c). Peak P3 contains the associated ribosome (Supplementary Fig. S1d), which is 75S in *Entamoeba* (Price et al., 1983), and small helical segments are also visible (Supplementary Fig. S1d), which might be fragments of helical ribosomal assemblies observed earlier (Morgan & Uzman, 1966). Further, peak P2 sample screening in cryo-EM conditions showed a uniform ribosomal distribution with optimum ice thickness (Supplementary Fig. S1e), suggesting the samples are ideal for high-resolution cryo-EM data collection.

### A rRNA triple helix constitutes the *E. histolytica* ribosomal large subunit

For ribosomal large subunit (LSU), the single particle reconstruction performed using RELION 3.1.4 has yielded an overall 2.8 Å resolution cryo-EM map from finally selected 378,061 particles (Fig. 1a, Supplementary Fig. S2 and Supplementary Table S1). The local resolution calculated using RELION showed the core of the ribosomal LSU has a resolution of up to 2.5 Å, whereas the surface has nearly 4.5 Å resolution. The cryo-EM map clearly shows the high-resolution features, such as resolved r-protein alpha helices and rRNA double helices (Fig. 1a, e, Supplementary movie 1). Remarkably, we could clearly see an rRNA triple helix in cryo-EM map, which is located near the peptide exit tunnel (PET) (Fig. 1a). The rRNA triple helix is composed of three helices, two parallel and one antiparallel helix. The three nucleotides are on the same plane. The nucleotide bases of the middle helix that is purine interact through Watson & Crick base pairing (–) with the nucleotide base of one of the helices and through Hoogsteen base pairing (•) with the nucleotide base of another helix. Four such nucleotides triplets U3326–A3414•U3311, U3325–A3425•U3312, U3324– A3426•U3313, and U3323–G3417•U3314 constitute the triple helix architecture (Fig. 1b, 1c, Supplementary movie 1). The position of the nucleotides are evident from its resolved cryo-EM density in our map (Fig. 1e, Supplementary movie 1).

**Figure 1.**
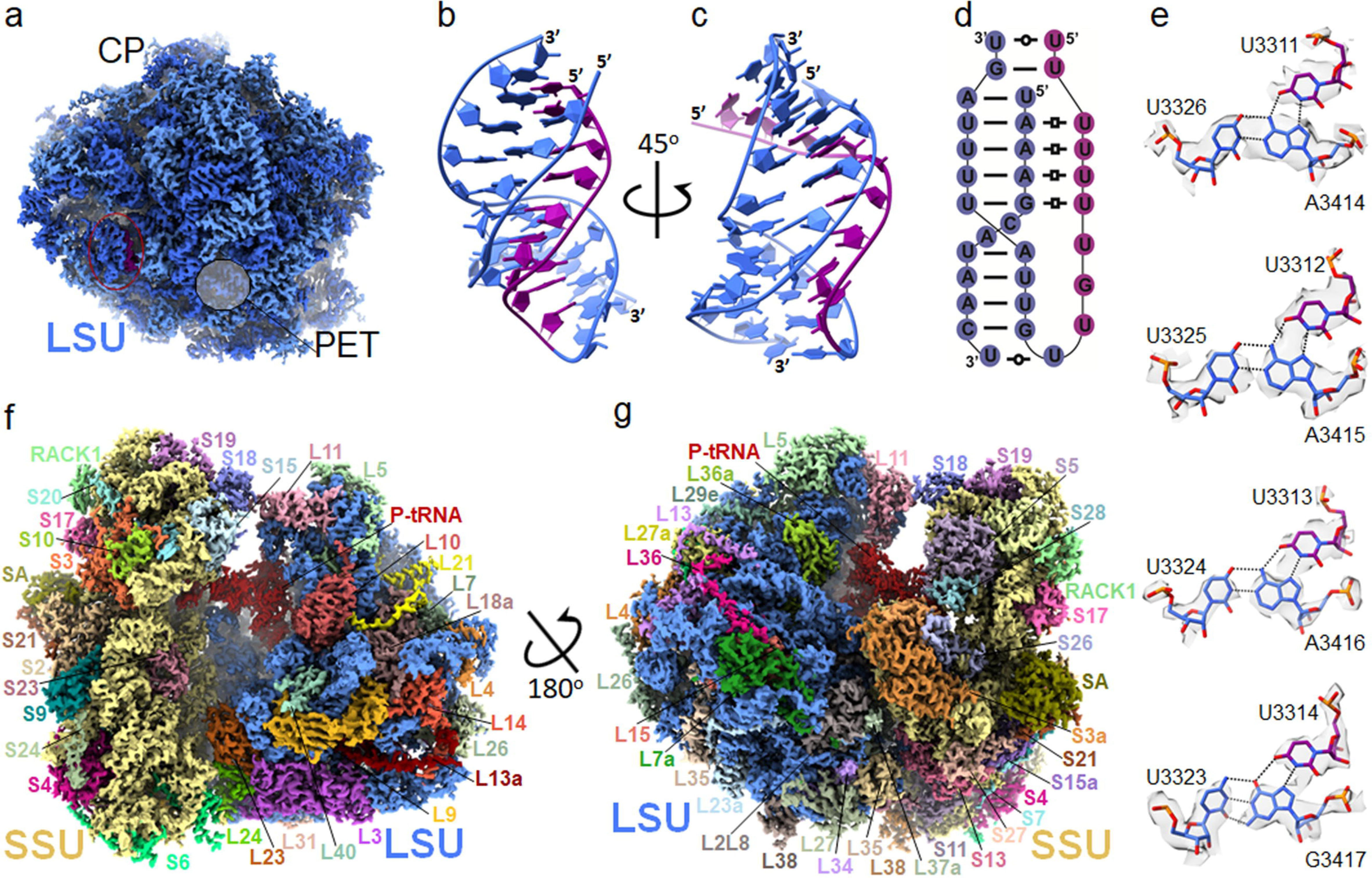
Cryo- EM structure of *E. histolytica* 53S and 7S ribosome. (a) The 2.8 Å resolution cryo- EM map of 53S ribosomal large subunit, rRNA (sky blue), r-proteins (royal blue), and third helix of rRNA triple helix (purple) are shown. Central perturbance (CP), large subunit (LSU), and peptide exit tunnel (PET) are labeled. A circle that is 50% transparent shows the location of PET in LSU. (b and c) rRNA triple helix, Watson and Crick pairing helices (sky blue), and third helix interacting to the major groove (purple), backbone in ribbon, and nucleotides in filled atoms are shown, and 5’ and 3’end (5’) are labeled. (b) and (c) are related by a rotation of 45° along the Y-axis. (d) A 2D diagram for triple helix color code and labeling is the same as for b and c. The **−** represents Watson and Crick base paired nucleotide and 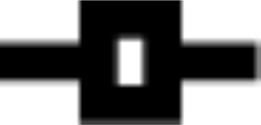 Hoogsten base paired nucleotides, and 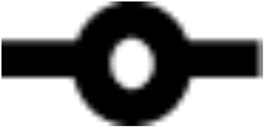 represents nucleotides in close proximity. (e) four base triples that constitute the triple helix are shown in the sticks, and cryo-EM density is shown on the surface with 85% transparency. (f, g) the 3.3 Å resolution cryo- EM 75S. rRNA of LSU (sky blue), SSU (khaki), and r-protein with different colors are shown. Ribosomal large subunit (LSU), small subunit (SSU), and r-protein are labeled. (f) and (g) related by a roughly 180° rotation along the Y-axis and 35° rotation along the Z-axis.

### The overall architecture of *E. histolytica* 75S ribosome

For the associated ribosome, which is 75S in *E. histolytica,* the cryo- EM single particle reconstruction yielded a 3.4 Å resolution map from 37,355 particles after particle polishing (Supplementary Fig. S3, Supplementary Table 1). The map’s resolution and overall quality improved further after multibody refinement with the final 3.3 Å resolution for both large and small subunits (Supplementary Fig. S3, Supplementary Table 1). The local resolution showed that the ribosome core has up to 3 Å resolution, whereas the periphery has nearly 5 Å resolution (Supplementary Fig. S3). Some of the high-resolution features similar to the large subunit cryo-EM map are also visible in the 75S ribosome map (Figures 1f, g and Supplementary Fig. S3), which enable us to build modified nucleotides (Fig. 2 and Supplementary Table S2). The 75S ribosome showed a P-site bound tRNA, suggesting that the tRNA is co-purified during ribosome purification (Fig. 1f, g).

**Figure 2.**
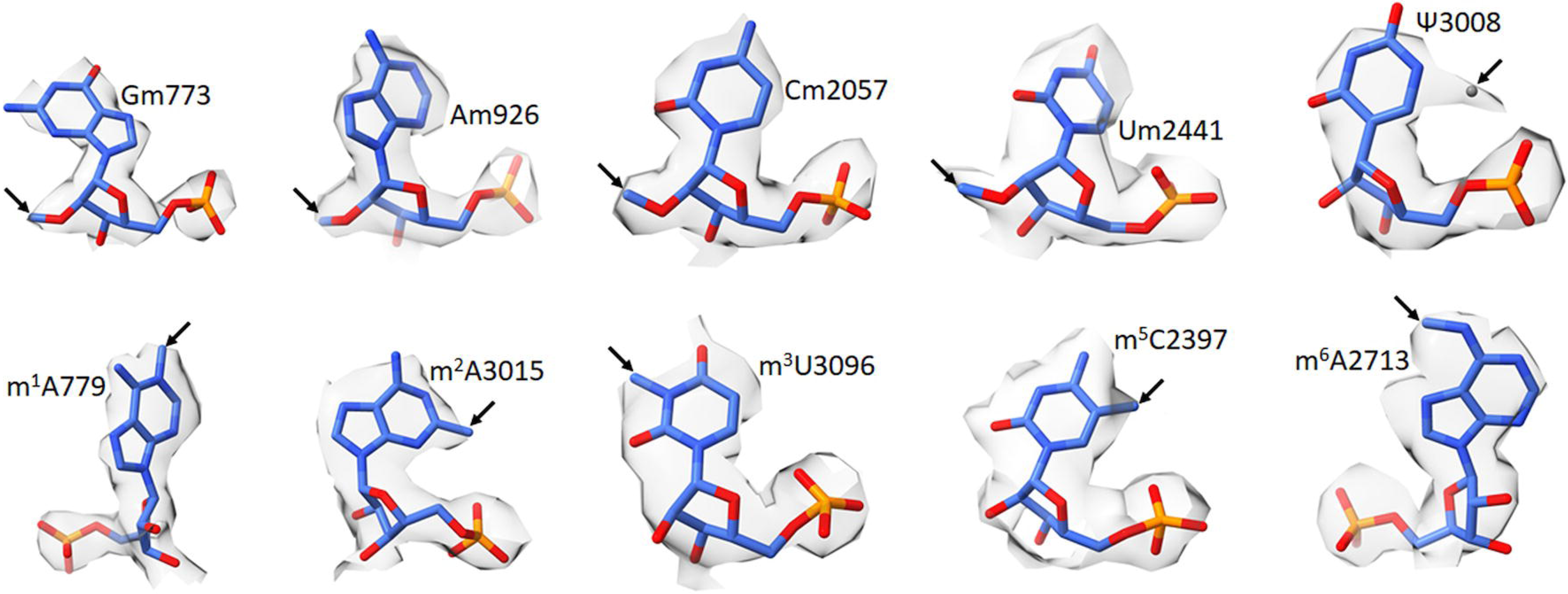
Modified nucleotides of *E. histolytica*. A representative of modified nucleotides modeled in cryo- EM maps are shown. The nucleotides in stick and cryo- EM density in surface view with 80% transparency is shown. Arrow makes the site of nucleotide modification. The grey • represents the modeled water molecule.

The cryo- EM maps of 53S and body 1 (LSU) from 75S clearly showed extra electron density at the 3’ end of 25S rRNA after modeling the annotated 25S rRNA sequence in the NCBI database (GenBank: BDEQ01000001.1). However, we could unambiguously build 41 extra nucleotides at the 3’ end and one nucleotide at the 5’ end (Supplementary Fig. S4). Therefore, we have reannotated the length of 25S rRNA to 3516 nucleotides, where 15 nucleotides out of an extra 56 nucleotides at the 3’ end of 25S rRNA remain disordered in our cryo- EM map (Supplementary Fig. S4). Approximately 92.46% of 25S (3251 out of 3516), 92.9% 5.8S (144 out of 155), and 100% 5S (117) rRNA were built in our cryo- EM map (Fig. 3 and Supplementary Fig. S5). Similarly, 74.88% of 17S (1458 out of 1947) nucleotides were built in cryo- EM map (Fig. 4 and Supplementary Fig. S6).

**Figure 3.**
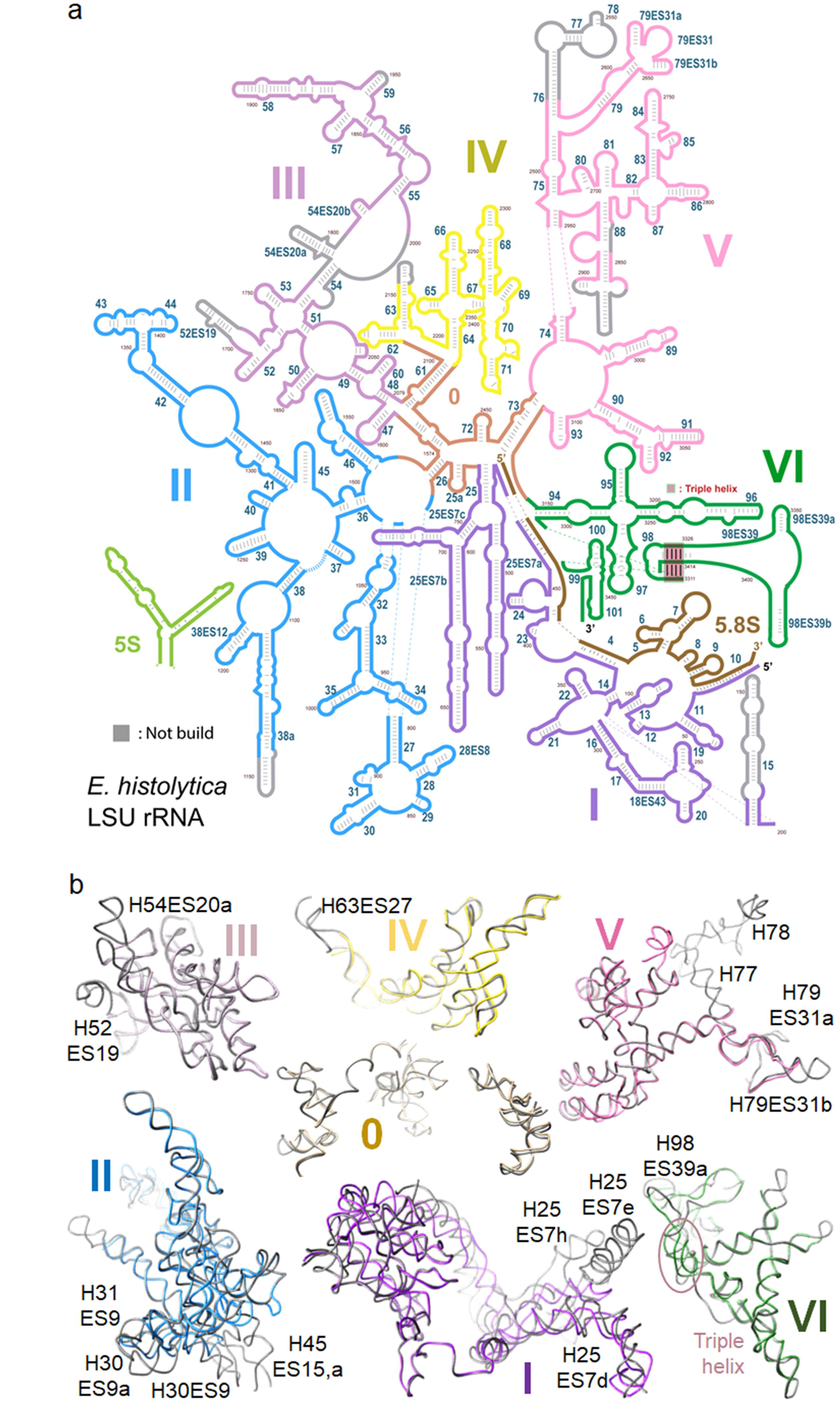
Secondary structure diagram of the *E. histolytica* LSU rRNA and its comparison with human counterparts. (a) 2D representation of the 25S, 5.8S and 5S rRNAs colored by domains. D0 (Sandy brown), DI (purple dark), DII (blue), DIII (purple light), DIV (yellow), DV (pink), DVI (green), 5.8S (sienna), and 5S (light green) are shown and labeled with a roman number. The rRNA that could not be modeled are shown in grey. Secondary structure were drawn by using R2DT tool from RNA Central and Adobe Illustrator and human templates for comparison was obtained from RiboVision suite (http://apollo.chemistry.gatech.edu/RiboVision). (b) Comparisons of individual *E. histolytica* 25S rRNA domains with the human 28S rRNA domains (grey) (PDB ID; 6QZP). The triple helix is circled and in an oval shape.

**Figure 4.**
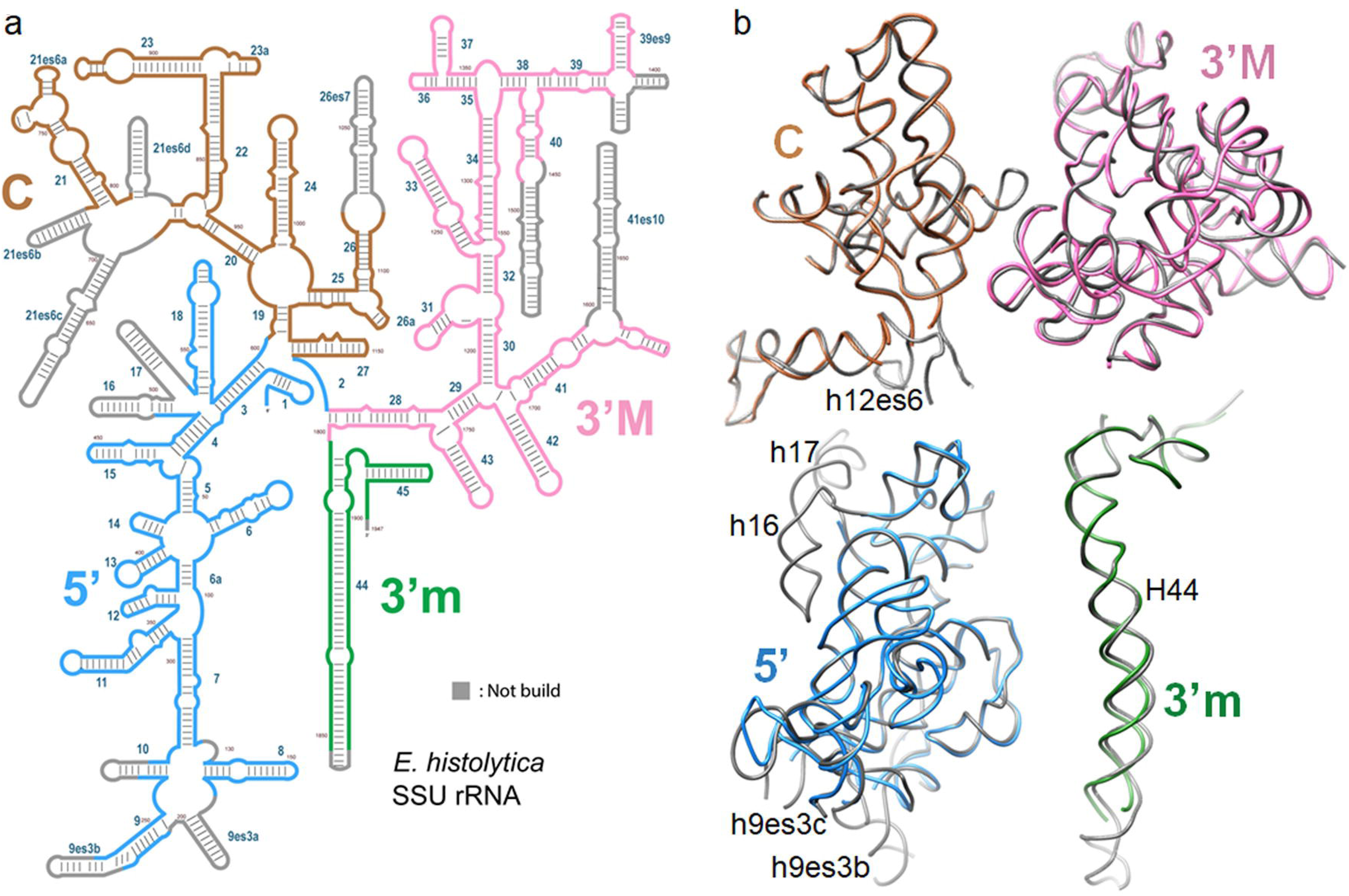
Secondary structure diagram of the *E. histolytica* SSU rRNA and its comparison with human counterparts. (a) 2D representation of the 17S rRNAs colored by domains. 5’ (blue), C (brown), 3’M (pink), 3m’ (green). The rRNA that could not be modeled are shown in grey. Secondary structure were drawn by using R2DT tool from RNA Central and Adobe Illustrator and human templates for comparison was obtained from the RiboVision suite (http://apollo.chemistry.gatech.edu/RiboVision). (b) Comparisons of individual *E. histolytica* 17S rRNA domains with the human 18S rRNA domains (grey) (PDB ID; 6QZP, Natchair) *et al*., 2017).

We were able to build 40 LSU and 32 SSU r-proteins (Fig. 1f, g) using 53S, body 1 (LSU) and body 2(SSU) from 75S. Also, our cryo- EM map allows us to assign the correct isoform for r-protein because of the clear density of the side chains (Supplementary Fig. S8, S9, S10). There were various discrepancies in the annotations of ribosomal protein where either they were not annotated or misannotated in the whole genome sequence available at NCBI with GANEBANK ID: BDEQ01000001.1. For LSU (Fig. 1f, g) protein L13 has six protein IDs(GAT95857, GAT97966, GAT97071, GAT99718, GAT92788 and GAT98013) and L13a (GAT97540) has one. Sequence alignment showed that (GAT95857, GAT97966, GAT97071, GAT99718) four out of six were identical to the L13a sequence, thus were misannotated as L13. 4 protein IDs, 2 each for eL10, 214aa long (GAT91935, GAT98189) and L10a, 210 aa long (GAT91778, GAT97776) are annotated in database but has similar sequence identical to L10 protein therefore, *Entamoeba* does not have L10a protein. Similarly, 3 protein IDs (GAT92445, GAT97165, GAT91560) mentioned as L44 show sequence similarity to L36a of humans and therefore reannotated as L36a. The genome database does not annotate Protein IDs EHI_141940, EHI_023220, which are L22 and L29, respectively. L29 is shorter (64 aa) and shares only 33% identity with its human counterpart (122 aa). We could build LSU r-proteins isoforms unambiguously in the cryo- EM map (supplementary Fig. S7, S8 and S9). Compared with human LSU r-proteins (PDB:6QZP), we found insertion and deletions in a few of the proteins, whereas proteins L10a, L28, and L41 were absent altogether. The cryo- EM density for some of the *E. histolytica* specific extensions were evident in the map (Fig. 1f, g), and we were able to build the extensions or insertions of L4, L9, L13, L18a, L26 and L31 (Supplementary Fig. S11). Intriguingly, L26 of *Entamoeba,* which has nearly 79 amino acid long insertion, overlaps the position of missing L28 protein. Whereas cryo- EM densities for some of *E. histolytica* specific deletion of r-proteins L3, L5, L6, L14, L18 and L29 were also missing in the cryo- EM map. Similarly, in SSU r-protein S8 (Protein ID: GAT98092) and RACK1(Protein ID: GAT91985) were not annotated and insertion in S21 and deletion in S17 were observed in comparison with human 40S ribosomal proteins. Most of the the SSU r-proteins are conserved, and some of the expansion regions of SSU proteins were not modeled as its cryo- EM density was absent, which may be because of its flexible nature.

### *E. histolytica* rRNA is reduced compared to its human counterparts Ribosomal large subunit rRNA

The LSU of *Entamoeba* consists of 25S, 5.8S and 5S rRNAs with 3516, 155 and 117, nucleotide length, respectively (Albach et al., 1984) (Fig. 3a, Supplementary Fig. S5). Over the period of evolution, the ribosome has gained various expansion segments (ESs) in rRNA to meet the increased functionality and regulatory requirement of the cell. This makes human LSU having an elaborate RNA having 28S, 5.8S, and 5S rRNAs with 5070, 157, 121 nucleotides length, respectively (Nachiar et al., 2017). On comparing both, the overall architecture of the rRNA core is conserved, whereas the surface helices are altered compared to their human counterparts (Fig 2b). The *E. histolytica* 25S rRNA can be subdivided into a familiar domain system for human 28S rRNA adapted from http://apollo.chemistry.gatech.edu/RibosomeGallery/index.html, which comprises of conserved core termed as domain 0 along with six other domains (D1-DVI) (Fig. 3a,b and Supplementary Fig S5).

Domain I in *E. histolytica,* composed of nucleotides 1 – 769 (769 nt. in length), is shorter than the human counterpart, having 1313 long nucleotides (Fig 3b, Supplementary Fig. S5). The shorter length can be attributed to either the absence of various expansion segments (H25ES7d, e, f, g, h) or their reduction, H25ES7a,7b, which are only 95 and 98 residues long as compared to human 246 and 242 residues, respectively (Fig. 3b). In addition to this the H25ES27c obtains a strikingly different conformation than its counterpart in human rRNA. Exceptionally, the H15 of 25S rRNA (87 nucleotides, disordered in our structure) is longer compared to 28S rRNA, the human equivalent (28 nucleotides) (Fig 3b, Supplementary Fig. S5).

Domain II in *E. histolytica* is from 794-1566, which is 257 nucleotides shorter than humans (1317-2346) and is almost similar except for the absence and reduction of a few expansion segments. Thus, altogether 30ES9a, 31ES9, 45ES15a, 45ES15 are the missing expansion segments, whereas H45, 30ES9, H31 are truncated in *Entamoeba* compared to humans. However, H46 has a 7 nucleotide long loop (1516-1523) in *Entamoeba,* which is absent in humans (Fig 3b, Supplementary Fig. S5).

Domain III, 492 nucleotides in length in *E. histolytica,* is longer than the human domain (2368-2827), comprising 459 residues. The extension can be ascribed to the expansion of H54ES20a (1769-1809, partially disordered), a 9 residue long loop (1600-1609 nt.), and the presence of an additional region after H55 (1998-2025) (Fig 3b, Supplementary Fig. S5).

Domain IV, comprising 298 nucleotides (2110-2408), is considerably reduced compared to 977 long human (2860-3837nt) counterparts. The domain is majorly composed of H63 and its expansion segments. The enormous difference is due to the absence of 668 nucleotide constituents of two major expansion segments 63ES27a and 63ES27b (Fig 3b, Supplementary Fig. S4, S5).

Domain V spans 2476-3121 residues (partially disordered) and contains almost similar nucleotides to humans (3903-4557). On the contrary, the domains show few prominent differences, which involve compact H78 with missing expansion segment 78ES30,30a (8 residues however, human is of 74) and a shorter 79ES31b. However, an EH-specific H88 expansion segment of 75 residues from 2841-2916(partially disordered) is not present in humans (Fig 3b, Supplementary Fig. S5).

Domain VI of *Entamoeba* is of 370 residues (3146-3516), shorter than the human 497 long domain (4572-5069). The reduction in length is due to reduced H98 and H101, where their constituent expansion segments 98ES39, 98ES39a, 98ES39b, and 101ES41, respectively, are compact (Fig 3b, Supplementary Fig. S5).

The 5.8S and 5S are mostly conserved with 155 and 117 nt in length, respectively, which are nearly same as its human counterparts (Fig 3b, Supplementary Fig. S5). The 5.8S and 5S in humans are 157 and 121 nt long, respectively.

Compared to human rRNA structure, we found that *E. histolytica* ribosomal RNA is compact, where most of the expansion segments are either truncated or missing. The ESs mostly evolved on the periphery of ribosome to meet other demands of the complex eukaryotic cells apart from protein synthesis, which includes transport from the site of biogenesis, adherence to the membrane of organelles, provide platform for co-post translational controls and site for binding of various ribosome biogenesis factors (Ramesh & Woolford, 2016), (Jeeninga et al., 1997), (Strunk & Karbstein, 2009); (Van Nues et al., 1997), (Fayet-Lebaron et al., 2009). The most widely studied ESs are H25ES7 and H63ES27. ES7 of LSU is truncated in *Entamoeba* where ES7d, e, and h are deleted, whereas a, b, and c are truncated, whereas whole ES27 was missing in EH. Removal of ES7 is lethal for fungi (Shankar et al., 2020). These expansion segments have various functions interacting with the ER membrane to provide anchorage to the organelle (Pfeffer et al., 2012), and provide a platform for the formation of enzyme bound ribosome complex involved in modifying N-terminal of emerging polypeptides in a co-translational manner. Its involvement has also been shown in the co-translational protein translocation in the 80S-Sec61 complex (Becker et al., 2009). Thus, *Entamoeba* translation fidelity may be compromised, or they might have another mechanism. However, it needs experimental evidence. Recently, it has been suggested that the H88 plays a role in phase 2 of the 60S ribosomal subunit biogenesis: the construction of central protuberance (CP). H88 and the L1 stalk provide two nucleation sites for the compaction and assembly of the RNAs (H80-H87) involved in forming CP and 5S RNP (Kater et al., 2020). Therefore, an *Entamoeba-specific* ES in H88 may be explored further regarding its involvement in CP reconstruction.

### Ribosomal small subunit rRNA

Similar to LSU 25S rRNA, the overall architecture of the *E. histolytica* SSU rRNA core is conserved (Fig. 1f, g). The peripheral helices are either missing or reduced in length compared to their human counterpart (Fig. 4, Supplementary Fig. S6). The SSU rRNA is 17S (Albach et al., 1984), and consists of 1947 nt. in contrast, its human counterpart is 1869 nt in length and its 18S. 17S rRNA is also made up of four domains, 5’ domain (5’), central (C) domain, 3’ major (3’M) domain and 3’ minor (3’m) domain (Fig. 4, Supplementary Fig. S6). *E. histolytica* 5’ domain is 602 nt (1-602nt) in length which is shorter compared to its human (658 nt) counterpart mainly because of missing expansion segments, h9es3c, b. whereas h16 and h17 remain disorder in our cryo- EM map (Fig. 4, Supplementary Fig. S6). *E. histolytica* C domain is 559 nt (603-1162 nt) in length, whereas it is 534 nt long in humans. But, still *Entamoeba* has missing h12es6. Some of the surface helices 21es6b,c,d and 25es7 are disordered in our cryo- EM map (Fig. 4, Supplementary Fig. S6). *E. histolytica* 3’M domain is 630 nt (1171-1801 nt) in length, exceptionally longer compared to human (497 nt) 3’M domain of SSU rRNA. The rRNA helices h39, h40 and h41 remain partially disordered in our cryo- EM map. *E. histolytica* 3’m domain is 145 nt (1802-1947nt) long and shorter than its human (169 nt) counterpart. The h44, which is the major helix in the 3’m domain, is shorter compared to the human h44 rRNA helix and partially disordered (Fig. 4, Supplementary Fig. S6). The 3’ M and 3’m domains constitute the decoding center on the SSU, where codon and anticodon interaction takes place during protein synthesis.

### *E. histolytica* rRNA and r-protein have co-evolved

Some of the r-proteins, such as L4, L9, L13, L26, and L31, possess prominent *E. histolytica* specific extensions or insertions in their sequences. These are mainly localized on the solvent side surface of the LSU (Fig. 5 and Supplementary Fig. S10). On the other hand, the LSU 25S rRNA is compact and reduced in size compared to human 28S rRNA (Fig. 3, Supplementary Fig. S5). The small size of 25S rRNA is mainly because some eukaryotic-specific expansion segments are missing (Fig. 3 and Supplementary Fig. S5). To stabilize the reduced size of rRNA, which has opted a different conformation for surface helices, *E. histolytica* has evolved with extensions, which contains mostly positively charged amino acid residues and adopt α helical secondary structure (Fig. 5). These helices sandwich between the rRNA helices, which are localized in the vicinity of the ribosomal surface (Fig. 5). For example, L4 extension sandwich between H25ES7b and H25ES7c, L9 insertion interacts with H42, L13 extension sandwich between H18ES43 and H25ES7a, similarly, the L26 extension sandwich between H45 and H46, and L31 interacts with the H94, H99 and H98ES39b, which constitutes the rRNA triple helix, an *E. histolytica* specific rRNA motif (Fig. 5). It suggests that the *E. histolytica* LSU rRNA helices and r-proteins have co-evolved to provide a compact and stable architecture, which might be crucial for its survival particularly in the cyst stage of its growth cycle.

**Figure 5.**
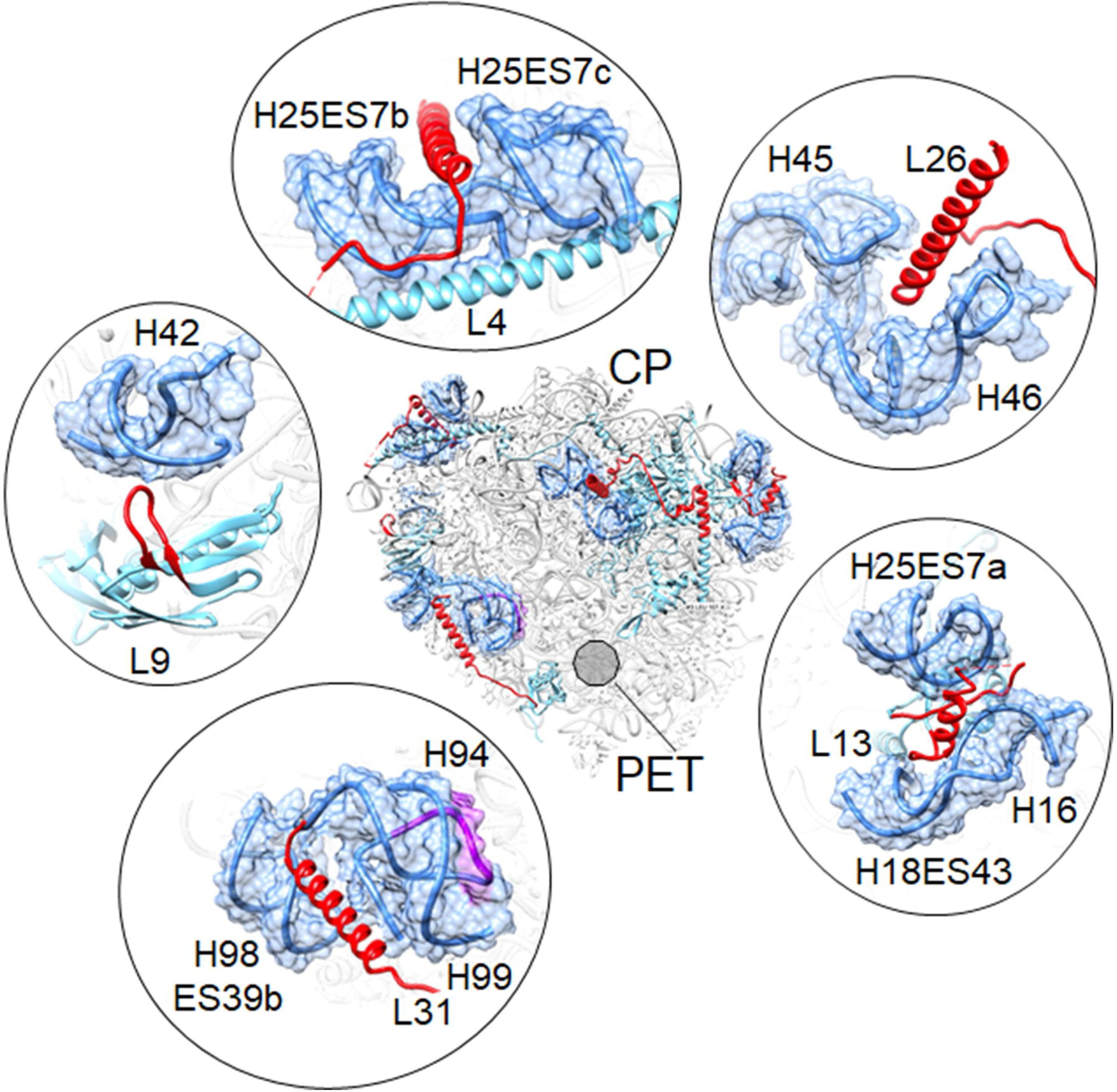
Coevolution of rRNA and r-proteins. The E. histolytica LSU is shown on the solvent side in the middle, and regions of rRNA and r-protein coevolution are zoomed and shown in the surrounding area in roughly matching orientation. The rRNA (blue) is shown in ribbon and surface view with 80% transparency. The r-proteins in the ribbon (light blue) and extension/insertion (red) are shown. Central perturbance (CP) and peptide exit tunnel (PET) are labeled. A circle that is 50% transparent shows the location of PET in LSU.

### RNA triple helix is a unique structure motif in RNA machinery

RNA triple helix, naturally found in many biological machineries, plays a vital role and provides catalytic scaffold, structural stability, metal binding site, etc. (Conrad, 2014; Brown, 2020). RNA triple helix is found in telomerase, an RNP enzyme that maintains the chromosome end, where an RNA triple helix is crucial for its catalytic activity (Nguyen et al., 2018). In spliceosome and Group II, introns RNA triple helix coordinate the catalytic Mg ^2+^ ions during splicing (Bertram et al., 2017; Qu et al., 2016). In riboswitches, RNA triple helix provides a ligand binding scaffold (Huang et al., 2017). RNA triple helix is also reported in human metastasis-associated lung adenocarcinoma transcript 1 (MALAT1), which is endowed with a blunt end triple helix (no 3’overhang), inhibits the rapid phase of nuclear RNA decay and also upregulates translation (Brown et al., 2014). Kaposi’s sarcoma-associated herpesvirus (KSHV) adopts a triple RNA helix motif in lytic phage and protects its genome from decay (Mitton-Fry et al., 2010).

Interestingly, we found a distinctly visible cryo- EM density for a triple helix in the *E. histolytica* LSU map (Fig. 1a-e, Supplementary movie 1). To the best of our knowledge, this is the first report of an RNA triple helix present in ribosomal RNA. The triple helix is present in DVI, towards the 3’ end of 25S rRNA, on the surface, solvent side, of the LSU near the PET (Fig. 1a, 3, Supplementary Fig. S5, S11, Supplementary movie 1). Compared with the human 28S rRNA, we observed that *E. histolytica* 3’ end containing domain six is reduced by 17 nucleotides. The human 28S rRNA 3’ position overlaps with the *E. histolytica* triple helix position on 25S rRNA (Supplementary Fig. S11). As the 3’ end is a target site for 3’ to 5’ exonucleases, it is speculative to say that *E. histolytica* has remodeled its 23S rRNA by shortening its length and adopting a different conformation of ’triple helix’ to protect its rRNA from human 3’ to 5’ exonucleases. It will most likely be similar to what was reported earlier for KSHV (Mitton-Fry et al., 2010). However, it needs experimental evidence.

### Paromomycin: a ribosome targeting antibiotic as an amoebicidal

Paromomycin (PAR), an aminoglycoside (AG), binds in the major groove of h44 in bacteria (Fig. 4, 6 Supplementary Fig. S6). The mechanism is well explained by (A. P. Carter et al., 2000). The ring 1 of PAR is involved in stacking interaction with G1491 and hydrogen bonding with A1408. PAR insertion into rRNA helix h44 flips out the two universally conserved adenines, A1492 and A1493, which otherwise need energy for flipping out. In flipped conformation, the nucleotides interact with cognate tRNA and facilitate the correct selection of tRNA in the A-site in normal protein synthesis (Schmeing & Ramakrishnan, 2009). In PAR bound state, these nucleotides, A1492 and A1493, remain permanently flipped out. Thus, it reduces energy requirements and facilitates the binding of both cognate and non-cognate tRNA, leading to error-prone translation. The other interactions with H69 of LSU and uS12 protein further stabilize PAR binding.

PAR is also used as an anti-parasitic agent to treat various protozoan diseases, and its mechanism of action is predominantly the same except for its interaction with eukaryotic specific proteins, eS30 and eL4. Mutations in both proteins show increased sensitivity to AGs (Dresios et al., 2006) (Fernández-Pevida et al., 2016)(Shalev-Benami et al., 2017). The PAR bound to ribosome from *L. donovani* structure (Shalev-Benami et al., 2017) suggests that the rRNA modifications govern the selectivity towards protozoan ribosome, and interestingly, *L. donovani* retains two of the bacterial specific m^4^Cm1402 and m^5^Cm1404 modifications (Fischer et al., 2015), which are located upstream to the PAR binding pocket. Together with the two conserved modifications m^6^ 2A at A1518 and A1519 in bacteria, which is also present in (and eukaryotes (Fig. 6). These modifications are located near the PAR binding pocket. It bridges the peripheral interactions like mRNA-tRNA codon-anticodon mini helix, decoding center helices h44 and h45 of SSU, and H69 of LSU. In our cryo- EM map, we could model bacterial specific modifications m^4^Cm1805, m^5^Cm1807, and the conserved modification m^6^2A at A1928 and A1929). In addition, G1491 of bacteria, in which ring1 of PAR stacks is conserved ’G1901’ in *E. histolytica* (Fig. 6). whereas it is mutated to A in both *Leishmania* and Humans. (Fig. 6). Thus, the *E. histolytica* PAR binding pocket is more similar to the bacterial than the eukaryotic.

**Figure 6.**
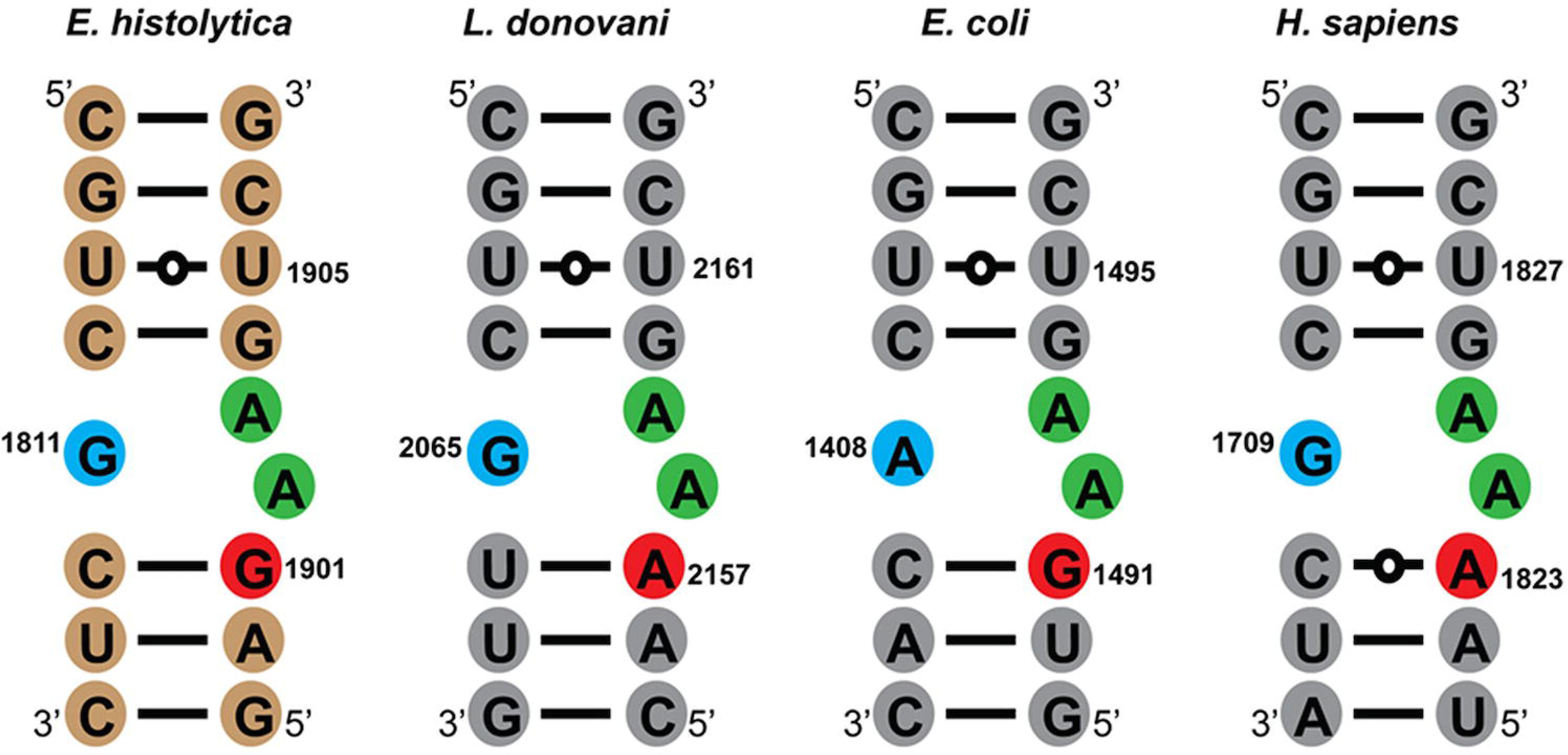
Helix 44, nucleotide of the paromomycin binding packet. A 2D diagram of helix 44 paromomycin binding pocket form *E. histolytica* (from this study), *L. donovani* (PDB ID; 6AZ1), *E. coli* (PDB ID; 5AFI), and *H. sapiens* (PDB ID; 6QZP) is shown. Universally conserved residues A1492 and A1493 (E. coli numbering) are shown with green background. Other crucial residues for paromomycin are shown with blue and red backgrounds. The chain polarity 5’ and 3’ are labeled.

## Methods

### Entamoeba histolytica cell growth

Trophozoites of *E. histolytica* strain HM1: IMSS cl-6 were grown axenically in TYI-S-33 medium at 35° C as described earlier by (Diamond et al., 1978), except where to get the sufficient amount of ribosomes culture was expanded into 75 cm^2^ cell culture, plug seal flask (NEST) with 250 ml complete media instead of glass culture tubes. In brief, TYI-S-33 media was prepared by mixing potassium phosphate dibasic 1.0 g, potassium phosphate monobasic 0.6 g, biosate peptone 30.0 g, glucose 10.0 g, sodium chloride 2.0 g, L-cysteine hydrochloride 1.0 g, ascorbic acid 1.0 g, and ferric ammonium citrate 22.8 g in 700 ml of double distilled water and pH was adjusted to 6.8 using 5N NaOH. The volume was made up to 900 ml (10 units) and filtered using filter assembly (corning 500 ml, filter system). The cells were grown in TYI-S-33 medium complemented with 15% adult bovine serum, 1X Diamond vitamin mix, 125 µl of 250 U/ml Benzylpenicillin, and 0.25 mg/ml Streptomycin per 90 ml of medium (complete media). After 60 hours, media was removed, and 1X Phosphate Buffer Saline (PBS pH 7.4) was added to the confluent culture, chilled on ice for 7 minutes, and then harvested by centrifugation at 1,200xg for 7 minutes at 4° C.

### Ribosome isolation and purification

Ribosome isolation was done similarly, as reported previously in (Kumar et al., 2024), with minor modifications. The *E. histolytica* cells were resuspended in lysis buffer (20 mM HEPES-sodium salt buffer pH 7.4, 40 mM KOAc, 10 mM Mg(OAc)_2_, 250 mM sucrose, 3 mM DTT, 1 mM PMSF, 1X Protease inhibitor cocktail (cOmplete, Roche), RNasin (Thermo,1:40 dilution ) and flash freeze in liquid nitrogen. The cell lysis was performed using Retsch Mixer Mill MM500 at 30 Hz, 1 minute/cycle, and a total of 8 cycles. 2 U/ml DNase I (Thermo Scientific) was added to the cell lysate and incubated on ice for 45 minutes. The cell debris was cleared by centrifugation at 18,626xg for 45 minutes at 4° C. The cleared cell lysate was layered on a 1.1 M sucrose cushion (20 mM HEPES-sodium salt buffer pH 7.4, 150 mM KOAc, 10 mM Mg(OAc)_2_, 1.1 M sucrose, 3 mM DTT) in equal volume, followed by ultra-centrifugation at 100,000g for 16 hours at 4° C (Hitachi P70AT2 rotor). The pellet was then resuspended in buffer A (20 mM HEPES-sodium salt buffer pH 7.4, 150 mM KOAc, 10 mM Mg(OAc)_2_, 3 Mm DTT) and centrifuged at 18,626xg 30 minutes, 4° C. The absorbance of the supernatant was measured at 260 nm, and 10 OD units were layered on a 10% - 50% sucrose gradient in gradient buffer (20 mM HEPES-sodium salt buffer pH 7.4, 150 mM KOAc, 10 mM Mg(OAc)_2_, 3 Mm DTT) prepared using Gradient Master from Biocomp Instruments. Gradients were centrifuged at 256,400xg for 8 hours at 4°C (Hitachi, P40ST rotor). Afterward, the gradients were fractionated using a Gilson Fractionator from BioComp Instruments (Supplementary Fig.1). The fractions were then analyzed using 1% agarose gel with 0.06% bleach (Supplementary Fig.1). The fractions corresponding to peaks were then pooled and concentrated using 100 kDa cut-off amicon from Merck Millipore, followed by buffer exchange with ribosome storage buffer (20 mM HEPES-sodium salt buffer pH 7.4, 100 mM KOAc, 10 mM Mg(OAc)_2_, 10 mM NH_4_OAc,3 mM DTT). The concentration of the sample was determined by absorbance at 260 nm, and ribosomes were stored at -80° C for future use.

### Electron microscopy

The 300 mesh Copper grids with a thin carbon layer on the top (CF300-Cu) from Electron Microscopy Science (EMS) were glow discharged for 50 seconds in a JEOL glow discharge unit. A 3 µl of sample from peak 1(Supplementary Fig.1a) of sucrose gradient fractionation was applied on the grid’s carbon side and washed three times with Mili Q water. The grid was stained with 1% uranyl acetate solution. The grids for peaks 2 & 3 Supplementary Fig.1a were prepared following a similar protocol. The grids were analyzed in 120 kVa, JEOL JEM1400 electron microscope at Advanced Technology Platform Centre (ATPC), Regional Centre for Biotechnology (RCB), Faridabad (Supplementary Fig.1b, c,d).

### Cryo- electron microscopy

A 3 µl of 53S ribosome (1µg/µl, peak 1) sample was applied to a quantifoil Cu 1.2/1.3 grids from Ted Pella, INC and plunge freeze using GATAN CP3 plunger after 3 second blotting at 4° C temperature and 80% humidity. The cryo- EM grid was mounted in a GATAN 626 cryo-holder and analyzed using a JEOL JEM2200FS microscope equipped with a GATAN K2 summit direct electron detector camera at ATPC, RCB Faridabad. At optimum ice thickness, the data was collected in a movie mode at 1.649 e-/Å^2^/frame with 30 frames per movie stack.

After initial cryo- EM condition optimization at ATPC, RCB, the final high-resolution data was collected at 300 kVa Titan Krios microscope equipped with Falcon 3 direct electron detector camera (ThermoFisher) at National Electron Cryo-Microscopy Facility, Bangalore Life Science Cluster (BLiSc), Bangalore. The data sets were collected in an electron integration movie mode. For the 53S ribosomal subunit, 5,724 movie stacks were collected with 25 movies per stack at 1.07 Å pixel with an electron dose of 1.34e/Å^2^/frame in electron integration mode using a Falcon 3 electron detector camera (Supplementary Table 1). For 75S ribosome, 3,470 movie stacks were collected in a similar setting as that of 53S ribosomal subunit data collection, with a similar electron dose of electron dose of 1.34 e/Å^2^/frame (Supplementary Table 1).

### Single particle reconstruction

Relion 3.1.4 (Zivanov et al., 2018) was used for single particle reconstruction of both 53S and 75S ribosome 3D reconstruction (Supplementary Fig 2,3 and Supplementary Table 1). The movie stacks were drift corrected using Relion’s own implementation, and single micrographs were generated. The micrographs were CTF corrected using CTFFIND4 (Rohou & Grigorieff, 2015). Initially, particles were picked manually, and 2D reference images were generated for automatic particle picking. 763,731 auto-picked particles for 53S ribosomal subunit were subjected to 2D classification, and the best classes with 554,832 particles were selected for 3D classification. A 60 Å lowpass filtered 60S ribosome cryo- EM map (EMDB ID 2660) from *Plasmodium falciparum* (Vong et al 2014), was used as a reference map. The three best 3D classes, consisting of 447,323 particles, were subjected to 3D refinement, which yielded a consensus map of 3.1 Å resolution after post-processing. The gold-standard FSC = 0.143 criterion (Rosenthal & Henderson, 2003) was used for resolution estimation through data processing. These 447,323 particles were subjected to CTF refinement and Bayesian polishing followed by 3D refinement and post-processing, which improved the overall map quality and resolution to 2.8 Å (Supplementary Fig 2). Further, the particles were subjected to a 3D classification into 3 classes without alignment, and one class with 378,061, which has homogeneous particles, was selected. The particles were 3D refined and post-processed to 2.7 Å resolution. Local resolution was estimated using ResMap (Kucukelbir et al., 2014), embedded in Relion.

Similar steps as that of 53S data processing were followed for the 75S ribosome refinement. 100,475 particles were auto-picked using 2D reference images generated after manual particle picking and 2D classification. The auto-picked particles were subjected to 2D classification, and the best classes consisting of 45,379 particles were selected for 3D refinement, and a consensus cryo- EM map was generated at 3.6 Å resolution. A lowpass filtered to 60Å cryo- EM map (EMDB ID 2660) *from Plasmodium falciparum* was used as a reference map in the 3D refinement. The particles were polished by CTF refinement and Bayesian polishing, which has further improved the overall resolution to 3.3 Å. A 3D classification into 3 classes without alignment was performed on the polished particles. The best class with 37,355 (82%) particles with tRNA bound at the P-site, subjected to 3D referment and post-processing, yielded a 3.3 Å resolution cryo- EM map. The resolution was estimated using the gold-standard FSC = 0.143 criterion (Rosenthal & Henderson, 2003). A multibody refinement (Nguyen et al., 2016) was performed to dissect the ribosomal inherent inter-subunit motion. The ribosomal large and small subunits were treated as body 1 and body 2, respectively. Body 1 was treated as fixed, and body 2 was refined by giving 10° rotation and 5 Å translation for multibody refinement. After post-processing the large subunit (body 1) and small subunit (body 2), the overall resolution remained the same, 3.3 Å. However, the map quality was improved further (Supplementary Fig 3). The final resolution was estimated using the gold-standard FSC = 0.143 criterion (Rosenthal & Henderson, 2003).

### Model building and structure analysis

rRNA and rProteins coordinates were built by coalescing both the approaches that is template based and de-novo model building. The atomic coordinates of *Toxoplasma gondii* (PDB ID: 5XXB) (Li et al., 2017) and *Homo sapiens* (PDB ID: 6QZP) (Natchiar et al., 2017) were used for template-based modeling where ribosomal rRNAs, 28S, 5S and 5.8S were rigid body docked into 53S cryo- EM using Chimera (Pettersen et al., 2004). By convention, 28S rRNA was divided into its 6 domains, and each domain was rigid body docked. The regions not fitting inside into the cryo- EM maps were deleted, and the nucleotide residues were mutated to the *E. histolytica* 25S rRNA, 5.8S, and 5S rRNA sequences using COOT v.0.9.3 (Casañal et al., 2020). The nucleotide sequences were manually built in the cryo- EM map using COOT for the remaining segments of rRNA. The r-protein modules were initially built using ModelAngelo (Jamali et al., 2022) without the FASTA sequence, followed by threading of the *Entamoeba* sequence on the carbon backbone by using PHYRE2 (Kelley et al., 2015). The iterative steps of manual model building, real space refinement, and geometry regularization were done in COOT, then phenix.real_space_refinement was used for the flexible refinement (Afonine et al., 2018).

For model building in the 75S 3.3 Å resolution map, the atomic coordinates of the 53S ribosomal subunit were rigid docked in cryo- EM map of the LSU (body 1) and followed by flexible refinement using phenix.real_space_refinement. The atomic coordinates of the 36S ribosomal subunit were built by following steps similar to that of the 53S model building in the 53S cryo- EM map described above. RNA modifications were manually modeled, and the quality of the density for the modifications is depicted in Fig: 2. The final coordinates of the 53S and 36S were rigid body docked in 75S cryo- EM map and followed by flexible refinement using phenix.real_space_refinement. The final model quality was checked using MolProbity (Prisant et al., 2020). The model building statistics are given in Supplementary Table 1. Figures were prepared in Chimera and ChimeraX (Goddard et al., 2018). The RNA 2D diagram was prepared using the R2DT tool (Sweeney et al., 2021)) at RNA Central using *Saccharomyces cerevisiae* and *Homo sapiens* as templates, RNAPDBee (Antczak et al., 2018)(Zok et al., 2018), (Antczak et al., 2014) and Adobe Illustrator.

## Data availability

Cryo- EM maps and corresponding coordinates are being submitted to EMDB and PDB databases, respectively.

## Acknowledgment

We acknowledge the National Electron Cryo-Microscopy facility at BLiSc, C-CAMP, Bangalore, a big thanks to Drs. Vinothkumar and Sucharita for cryo- EM data collection. We acknowledge the Advanced Technology Platform Centre at the Regional Centre for Biotechnology, Faridabad, and thanks Dr. Reena and Mr. Madhava for help in the initial cryo- EM sample screening. We acknowledge the computing resource at the Indian Biological Data Centre (IBDC), Faridabad. We thank all PSK lab members for proofreading the manuscript and providing valuable insights. Faridabad. SS acknowledges a fellowship from DBT. SM acknowledges Fellowship from UGC and ICMR. This work is supported by a research grant, BT/PR45101/DRUG/134/121/2022, awarded to S.G. and P.S.K. from the Department of Biotechnology, India.

## Author contributions

P.S.K. connived the research project. S.M. grew the *E. histolytica* pellet. S.S. purified and characterized the ribosomes. P.S.K. and S.S. performed the single particle reconstruction, model building, and structure analysis. P.S.K. and S.S. wrote the manuscript with input from S.M. and S.G..

