## Supplementary Figures for "Cryo- EM structure of ribosome from pathogenic protozoa *Entamoeba histolytica,* reveals unique features of its architecture"

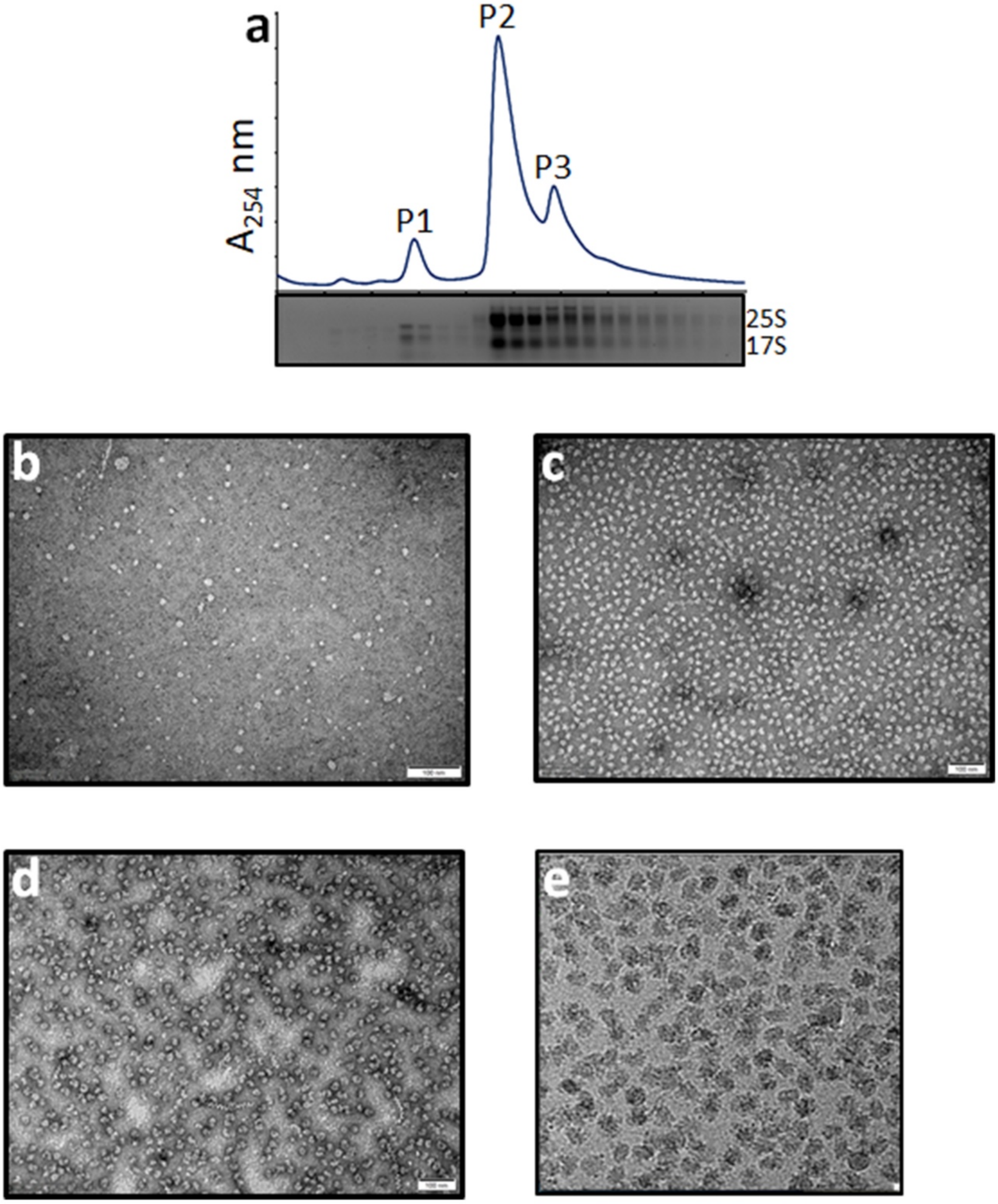


**Supplementary Figure S1. Characterization of E. histolytica ribosomes.** (a) Sucrose density gradient centrifugation (SDGC) and fractionation profile using 10% to 50% sucrose gradient and absorbance at 254 nm P1 (peak1), P2 (peak2), and P3 (peak3) are labeled. The fraction analyzed using 1% agarose with 0.06% bleach is shown below the SDGC profile; bands corresponding to 25S and 17S rRNA are labeled. (b) a representative 2D micrograph of the peak1 (P1) fraction. (c) a representative 2D micrograph of the peak2 (P2) fraction. (d) a representative 2D micrograph of the peak3 (P3) fraction. the negative staining 2D micrographs, b – d, were stained with 1% uranyl acetate. (e) a representative Cryo- EM 2D micrograph of P2 fraction.

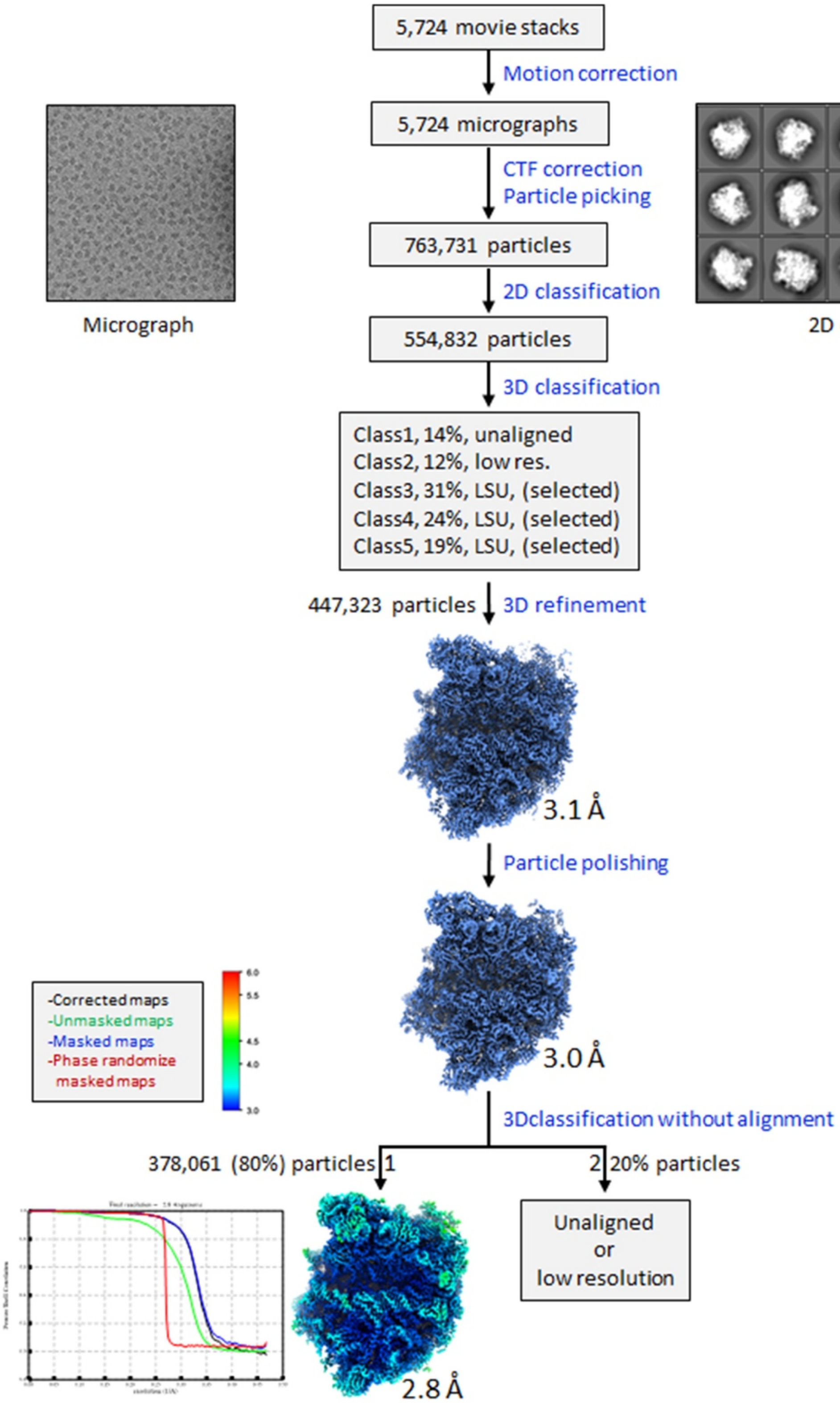


**Supplementary Figure S2. Summary of ribosomal large subunit cryo- EM single particle reconstruction.** A representative 2D micrograph and 2D class average are shown at top left and top right, respectively. Data statistics, no. of movies, no of micrographs, and no of particles used in image processing are listed, 3D maps after initial refinement, particle polishing, and final map with local resolution are shown. A Fourier shall correlation for the final cryo-EM map after postprocessing and masking are shown at the bottom left.

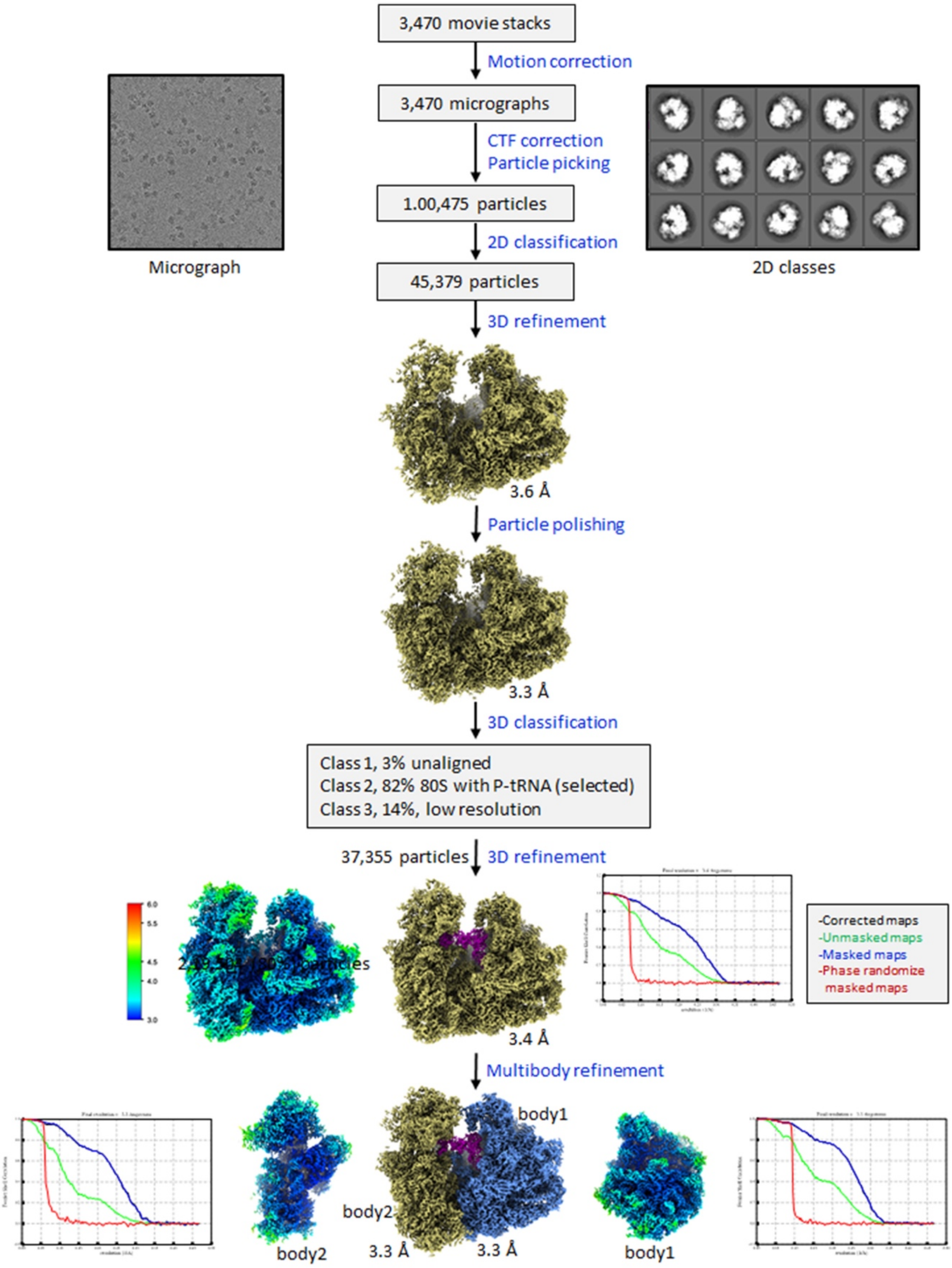


**Supplementary Figure S3. Summary of associated ribosome cryo- EM single particle reconstruction.** A representative 2D micrograph and 2D class average are shown at top left and top right, respectively. Data statistics, no. of movies, no of micrographs, and no of particles used in image processing are listed, 3D maps after initial refinement, particle polishing, multibody refinement and final map for body1 and body2 with local resolution are shown. A Fourier shall correlation for the final cryo-EM maps after postprocessing and masking also shown.

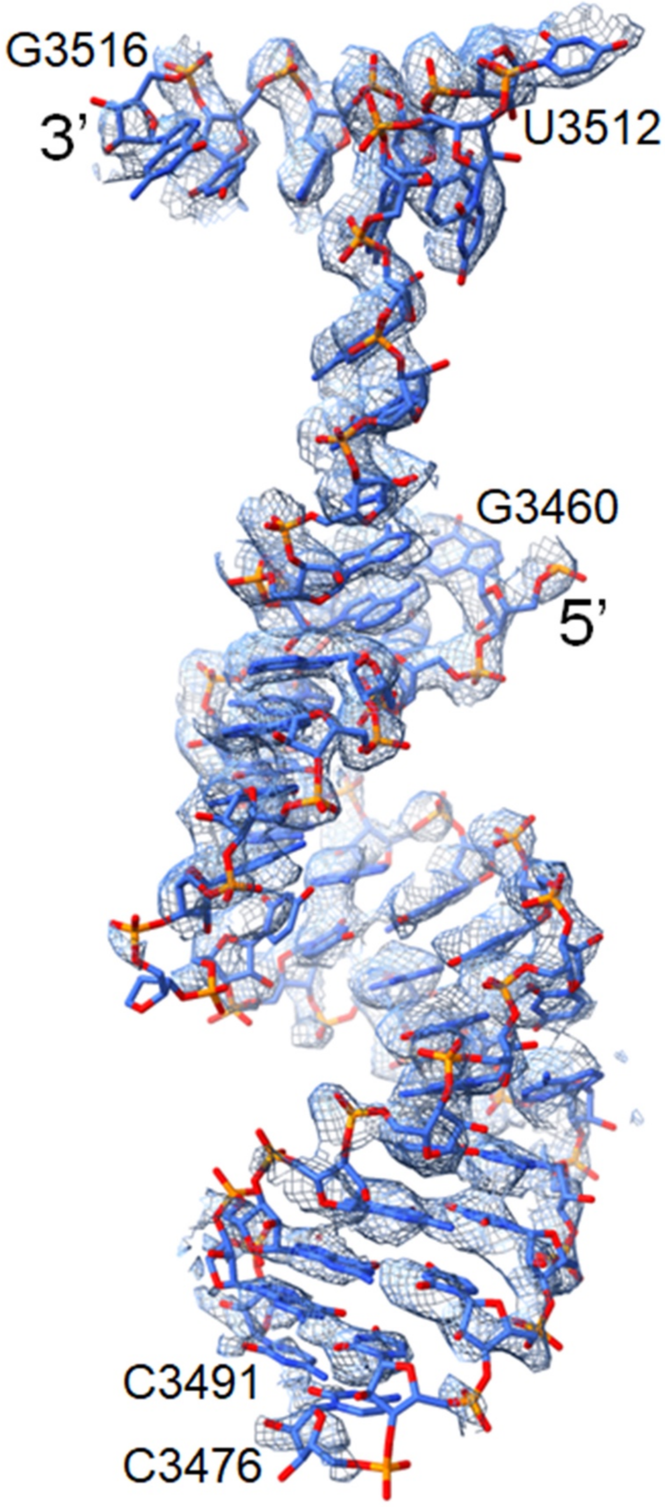


**Supplementary Figure 4. Ribosomal large subunit rRNA 3' end modeling.** The ribosomal large subunit 3' end from nucleotide 3460 to nucleotide 3516, modeled in cryo-EM map using coot, is shown. Regions after nucleotide 3476 and before nucleotide 3491 remain disordered. Cryo-EM electron density is shown in blue mesh with 80% transparency, and nucleotides are shown in stick representation.

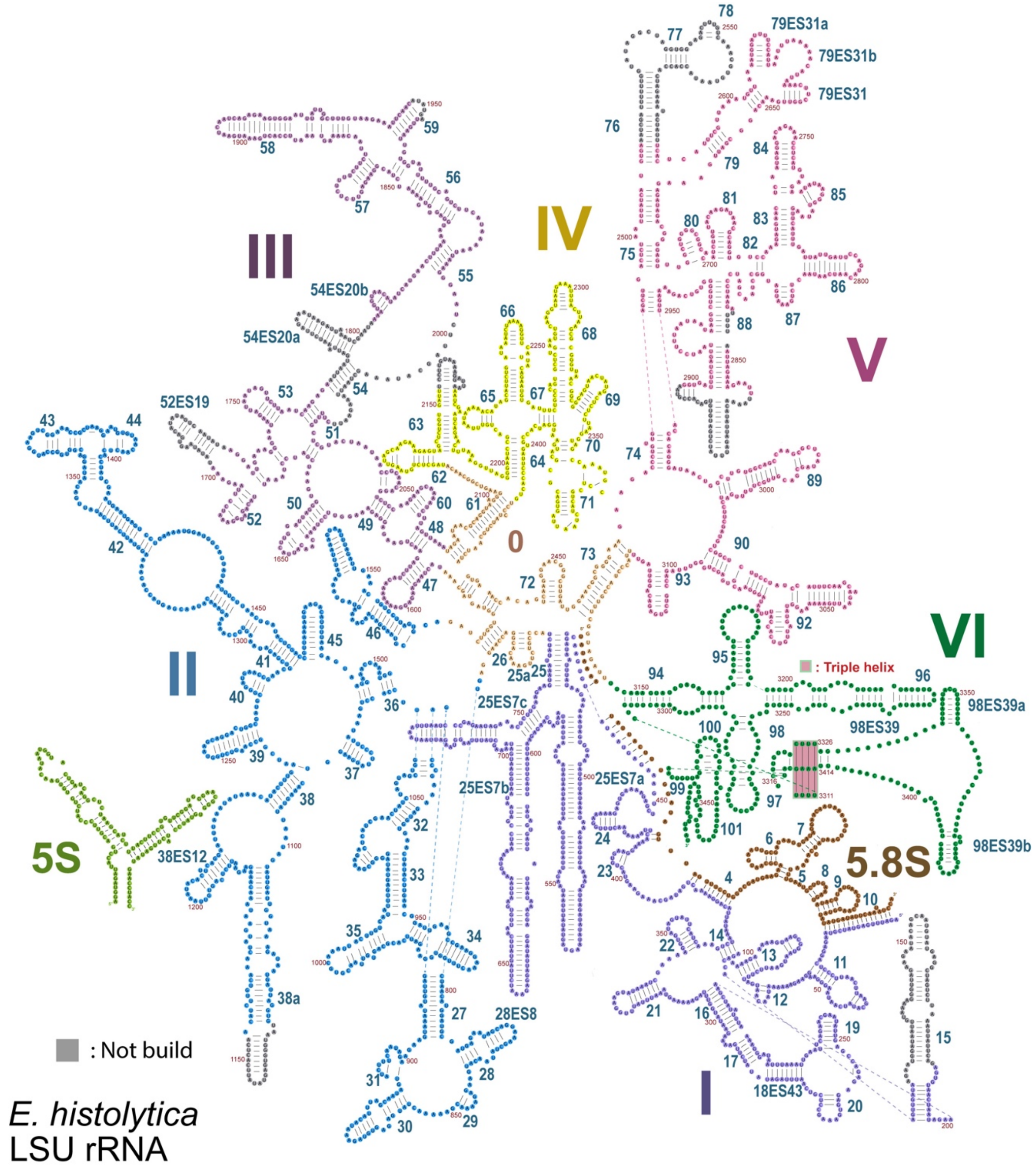


**Supplementary Figure 5. 2D representation of ribosomal large subunit rRNA.** Nucleotides in single letter code, residues of helices –, rRNA domains 0 (khaki), I (magenta), II (blue), III (Purple), IV yellow, V (red) and VI (green) are shown. 5.8S rRNA (Brown), triple helix (red background with 50% transparency), and 5S (light green) bottom left are shown. Nucleotides in grey are not modeled in cryo- EM map.

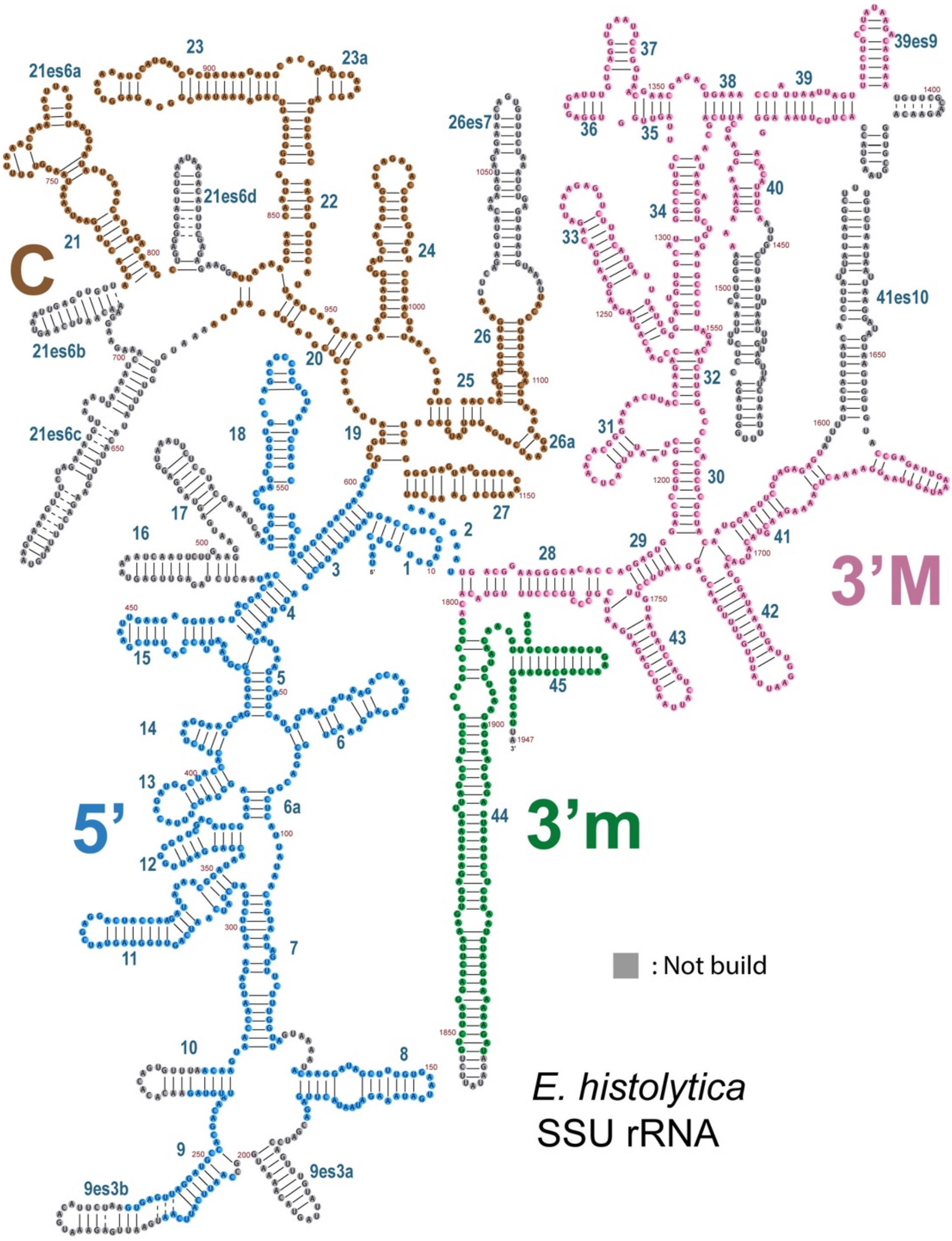


**Supplementary Fig. 6 2D representation of ribosomal small subunit rRNA.** Nucleotides in single letter code, residues of helices –, rRNA, 5' domain (sky blue), C domain (brown), 3'M domain (magenta) and 3' m (green) are shown. Nucleotides in grey are not modeled in cryo-EM map.

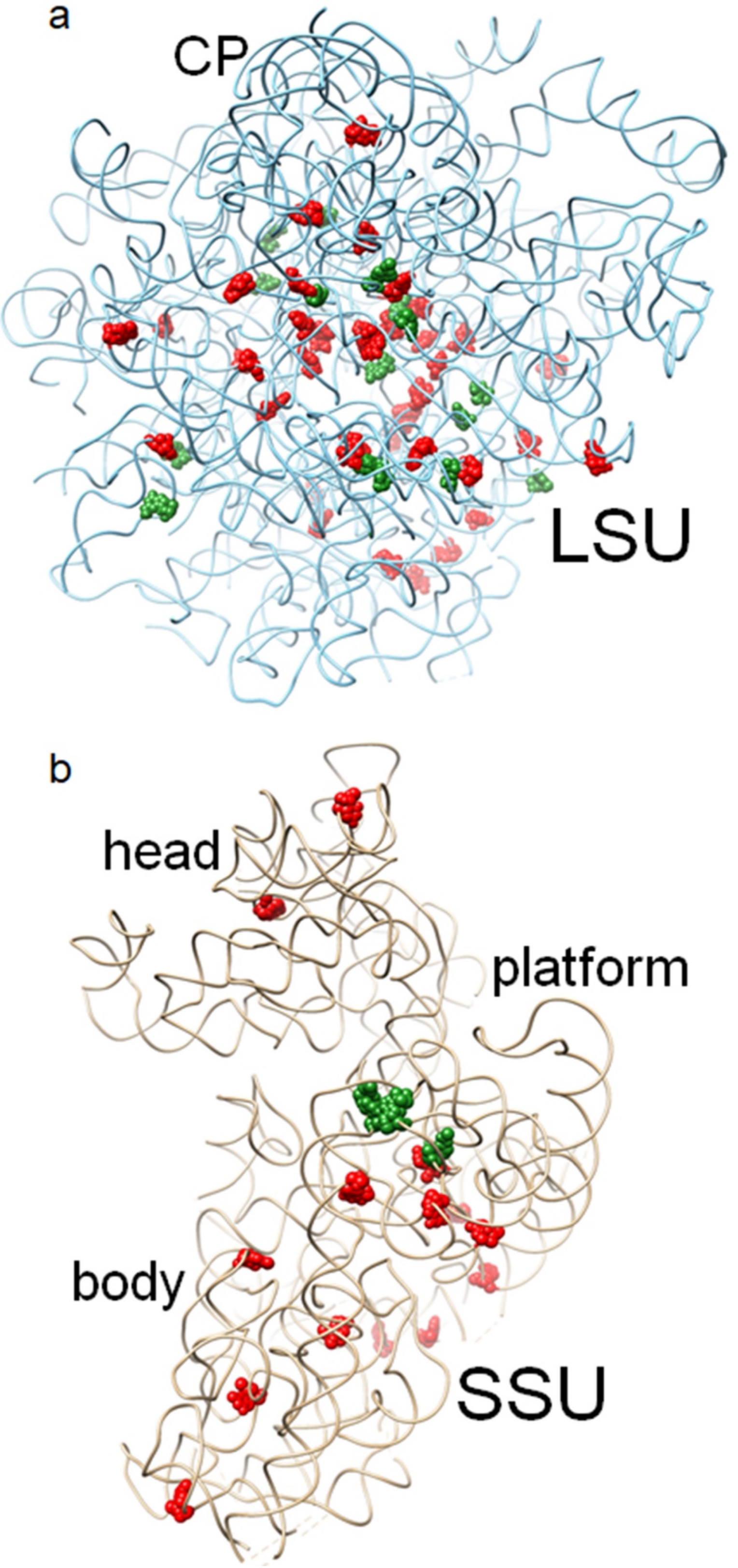


**Supplementary Fig. 7. rRNA base modifications.** (a) LSU rRNA (blue), (b) SSU rRNA (khaki). rRNA shown in ribbon, modified nucleotides are shown in the atom sphere, sugar modification (red), and base modification (green).

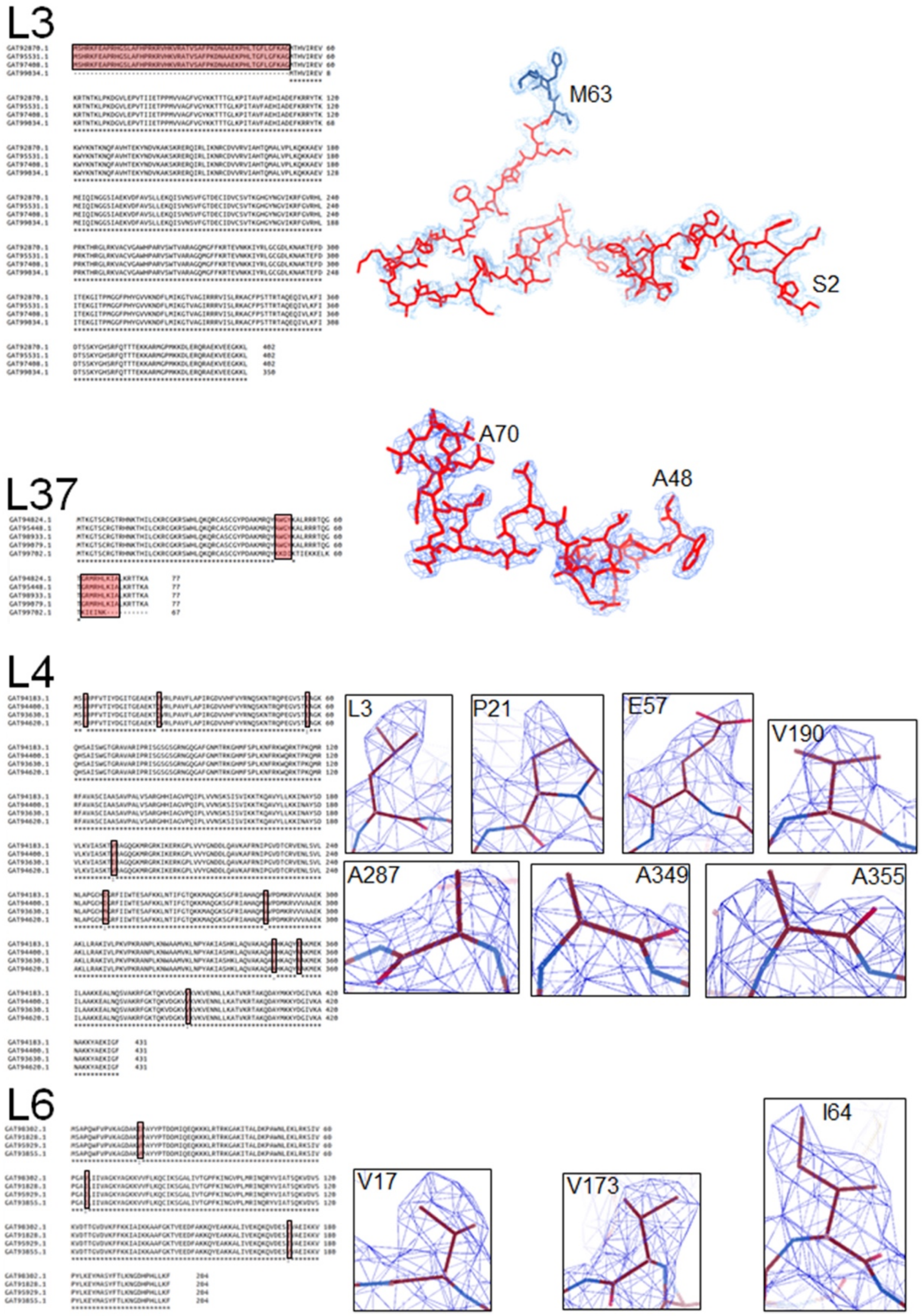


**Supplementary Fig. 8. r- protein L3, L37, L4 and L6 isoform modeling.** (Left panel) sequence alignment of all the isoforms of r-protein. The difference in the sequence is boxed and highlighted with a red background. (right panel) cryo- EM density in mesh and modeled amino acid residue in stick are shown.

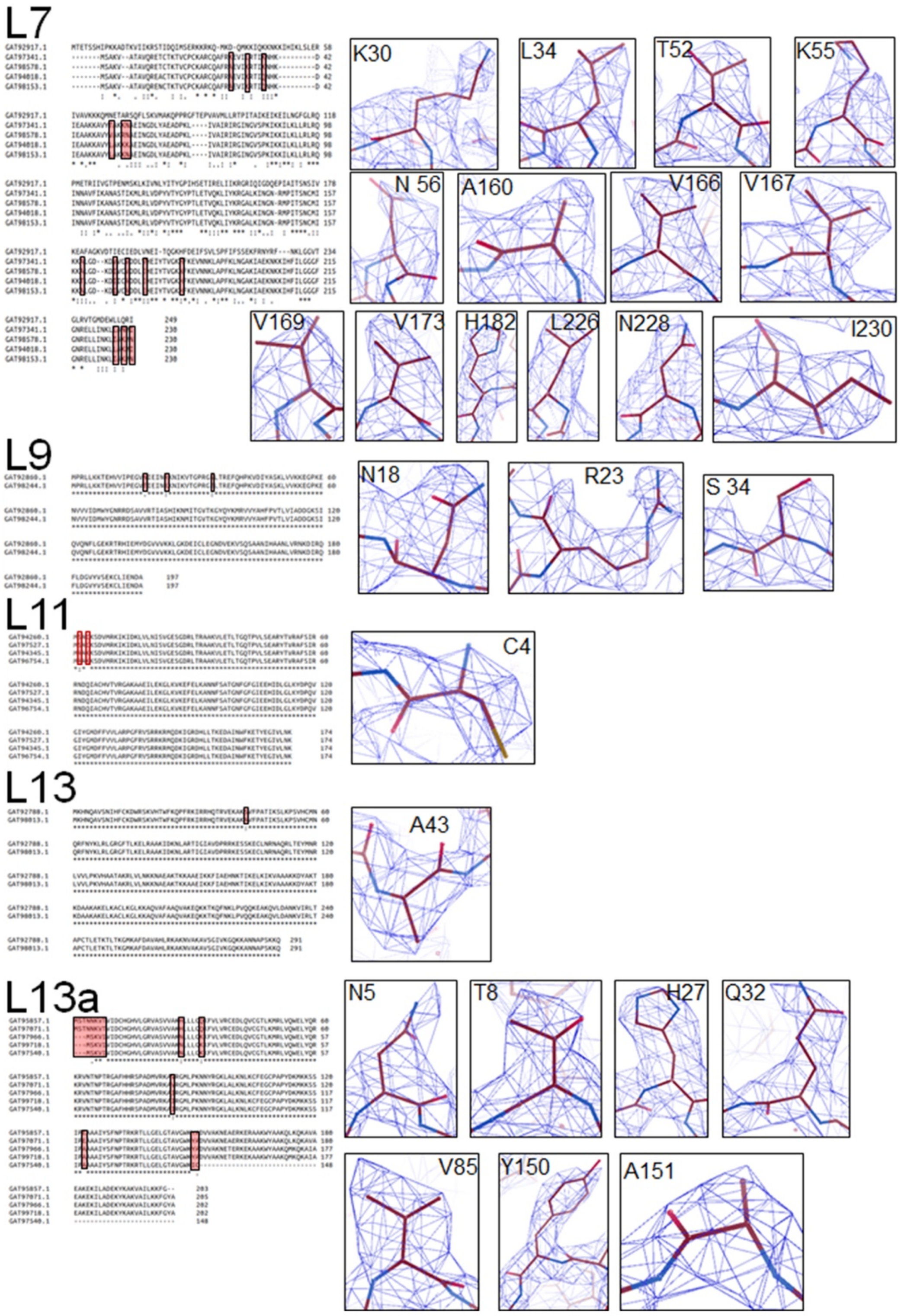


**Supplementary Fig. S9. r- protein L7, L9, L11, L13 and L13a isoform modeling.** (Left panel) sequence alignment of all the isoforms of r-protein. The difference in the sequence is boxed and highlighted with a red background. (right panel) cryo- EM density in mesh and modeled amino acid residue in stick are shown.

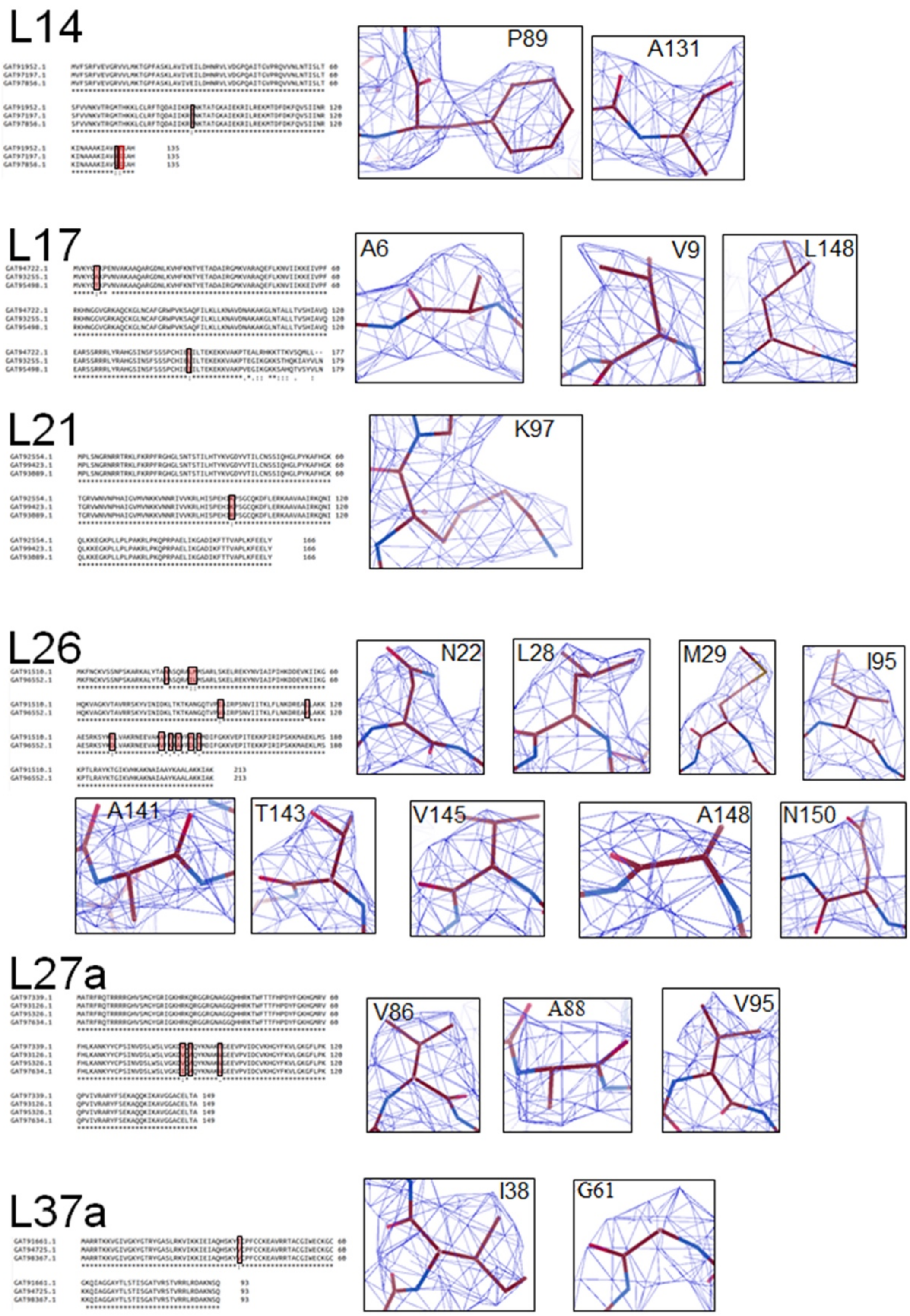


**Supplementary Fig. S10. r- protein L14, L17, L21, L26, L27a and L37a isoform modeling.** (Left panel) sequence alignment of all the isoforms of r-protein. The difference in the sequence is boxed and highlighted with a red background. (right panel) cryo- EM density in mesh and modeled amino acid residue in stick are shown.

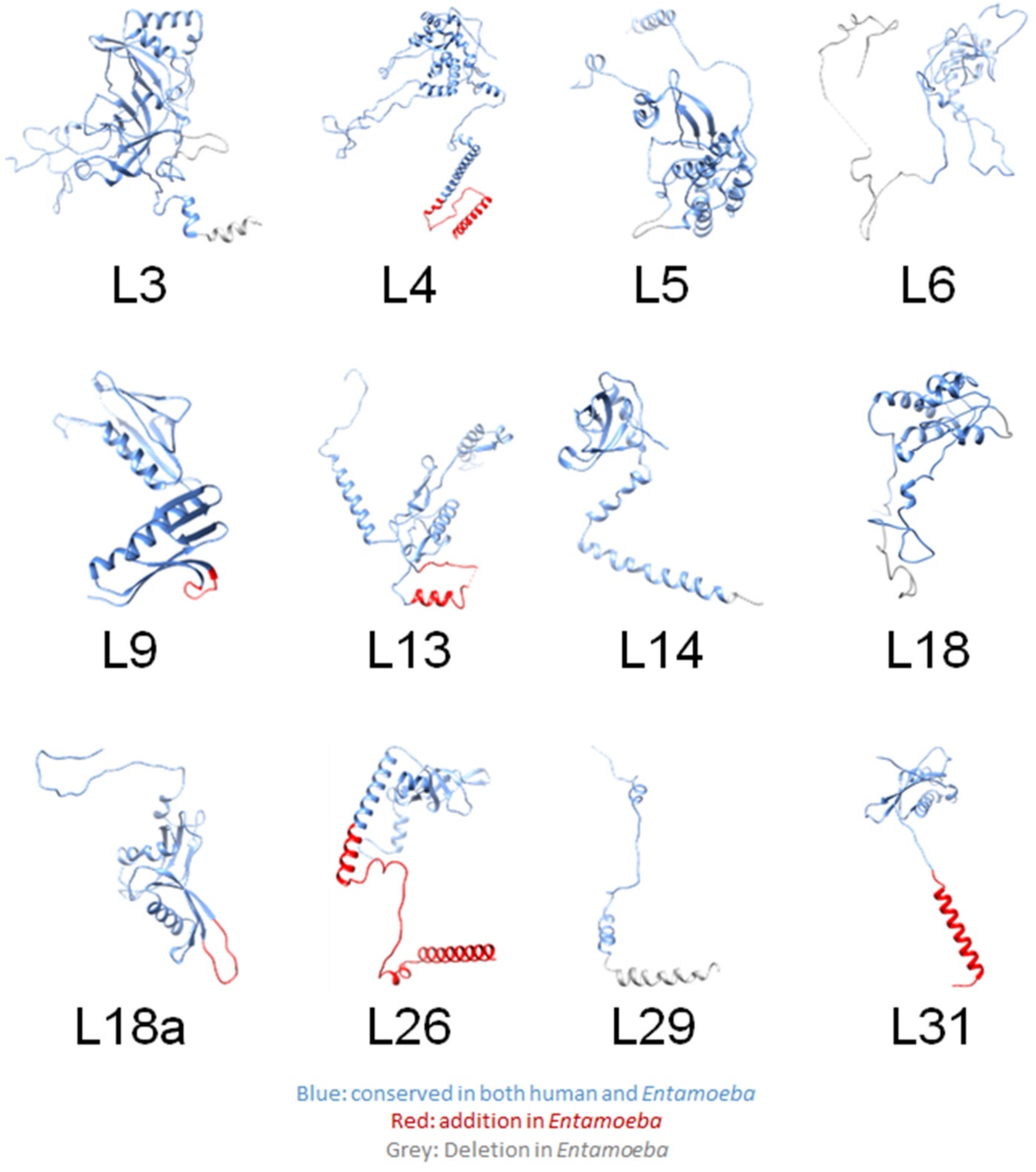


**Supplementary Fig. S11. Altered LSU r-proteins.** The r-protein is shown in the ribbon diagram. Conserved segments between the *Entamoeba* and human (blue), unique to the *Entamoeba* (red) and missing in the *Entamoeba* (grey), are shown.

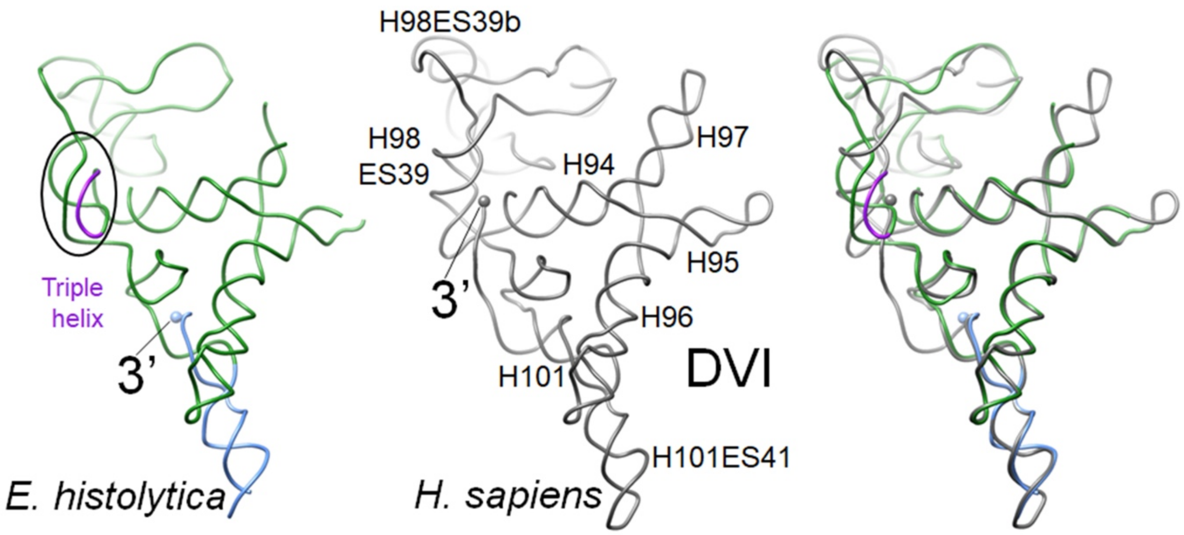


**Supplementary Fig. S12. Comparative analysis of LSU rRNA domain VI. (**left panel) 25S rRNA domain VI from *E. histolytica* (green), one the helix of the triple helix (magenta), extra nucleotides modeled based on cryo- EM density (sky blue) are shown. (middle panel) the 28S rRNA, domain VI from *H. sapiens* (grey) is shown and major helices are labelled. (right panel) the superimposition of domain VI from *E. histolytica* and *H. sapiens* are shown. For clarity, helices are labeled in the middle panel only.
