## Supplementary Table1 for "Cryo- EM structure of ribosome from pathogenic protozoa *Entamoeba histolytica,* reveals unique features of its architecture"

**Supplementary Table 1. Cryo-EM data collection, refinement, and validation statistics**

| | LSUS (EMDB-yyyyy PDB-XXXX | 75S EMDB-yyyyy PDB-XXXX | LSU (body 1) EMDB-yyyyy PDB-XXXX | SSU(body2) EMDB-yyyy PDB-XXXX |
|---|---|---|---|---|
| **Data collection and processing** | | | | |
| Magnification | 70,000 | 70,000 | | |
| Voltage (kV) | 300 | 300 | | |
| Electron exposure (e–/Å$^2$) | 1.34 | 1.34 | | |
| Defocus range (μm) | -0.5 to -3 | -0.5 to -3 | | |
| Pixel size (Å) | 1.07 | 1.07 | | |
| Symmetry imposed | C1 | C1 | | |
| Initial particle images (no.) | 763,731 | 100,475 | | |
| Final particle images (no.) | 378,061 | 37,355 | | |
| Map resolution (Å) | 2.8 | 3.3 | | |
| FSC threshold | 0.143 | 0.143 | | |
| Map resolution range (Å) | 2.5 to 5.0 | 3.0 to 5.5 | | |
| **Refinement** | | | | |
| Initial model used (PDB code) | 5XXB & 6qZP | XXXX | XXXX | 5XXB & 6qZP |
| Map sharpening *B* factor (Å$^2$) | -70 | -80 | -75 | -75 |
| Model composition | | | | |
| RNA nucleotides | 3402 | 11577 | 3402 | 1458 |
| Protein residues | 6175 | 10384 | 6175 | 4209 |
| Validation | | | | |
| Clashscore | 11.68 | | 11.21 | 16.35 |
| Poor rotamers (%) | 2.53 | | 0.15 | 0.49 |
| Ramachandran plot | | | | |
| Favored (%) | 93 | 91 | 92 | 92 |
| Allowed (%) | 6 | 7 | 7 | 7 |
| Disallowed (%) | 1 | 2 | 1 | 1 |
