## Supplementary Table2 for "Cryo- EM structure of ribosome from pathogenic protozoa *Entamoeba histolytica,* reveals unique features of its architecture"

**Supplementary Table 2. List of rRNA nucleotide modification**
**LSU modifications**

| Chain | Sl. no | Nucleotide no. | Nucleotide | Modification type |
|---|---|---|---|---|
| Chain 1 | 1 | 47 | A | A2M |
| Chain 1 | 2 | 59 | A | A2M |
| Chain 1 | 3 | 424 | A | A2M |
| Chain 1 | 4 | 425 | C | OMC |
| Chain 1 | 5 | 773 | G | OMG |
| Chain 1 | 6 | 779 | A | 1MA |
| Chain 1 | 7 | 780 | A | A2M |
| Chain 1 | 8 | 783 | A | A2M |
| Chain 1 | 9 | 798 | U | OMU |
| Chain 1 | 10 | 910 | A | 2MA |
| Chain 1 | 11 | 924 | G | OMG |
| Chain 1 | 12 | 926 | A | A2M |
| Chain 1 | 13 | 1027 | G | OMG |
| Chain 1 | 14 | 1079 | U | PSU |
| Chain 1 | 15 | 1091 | A | 2MA |
| Chain 1 | 16 | 1180 | A | A2M |
| Chain 1 | 17 | 1258 | A | A2M |
| Chain 1 | 18 | 1583 | A | A2M |
| Chain 1 | 19 | 1584 | G | OMG |
| Chain 1 | 20 | 1649 | G | OMG |
| Chain 1 | 21 | 1682 | A | 2MA |
| Chain 1 | 22 | 1751 | A | A2M |
| Chain 1 | 23 | 2057 | C | OMC |
| Chain 1 | 24 | 2063 | A | A2M |
| Chain 1 | 25 | 2217 | A | A2M |
| Chain 1 | 26 | 2295 | A | A2M |
| Chain 1 | 27 | 2354 | C | 5MC |
| Chain 1 | 28 | 2357 | A | A2M |
| Chain 1 | 29 | 2364 | G | OMG |
| Chain 1 | 30 | 2372 | A | A2M |
| Chain 1 | 31 | 2373 | U | 5MU |
| Chain 1 | 32 | 2397 | C | 5MC |
| Chain 1 | 33 | 2413 | C | OMC |
| Chain 1 | 34 | 2439 | A | A2M |
| Chain 1 | 35 | 2441 | C | OMC |
| Chain 1 | 36 | 2471 | G | OMG |
| Chain 1 | 37 | 2486 | U | OMU |
| Chain 1 | 38 | 2497 | U | OMU |
| Chain 1 | 39 | 2595 | G | 2MG |
| Chain 1 | 40 | 2665 | A | A2M |
| Chain 1 | 41 | 2689 | G | OMG |
| Chain 1 | 42 | 2711 | U | OMU |
| Chain 1 | 43 | 2713 | A | 6MZ |
| Chain 1 | 44 | 2794 | U | OMU |
| Chain 1 | 45 | 2936 | G | OMG |
| Chain 1 | 46 | 2941 | C | OMC |
| Chain 1 | 47 | 2958 | G | OMG |
| Chain 1 | 48 | 3008 | U | PSU |

| | | | | |
|---|---|---|---|---|
| Chain 1 | 49 | 3013 | C | 5MC |
| Chain 1 | 50 | 3015 | A | 2MA |
| Chain 1 | 51 | 3023 | U | 5MU |
| Chain 1 | 52 | 3025 | U | 5MU |
| Chain 1 | 53 | 3089 | A | A2M |
| Chain 1 | 54 | 3102 | C | OMC |
| Chain 1 | 55 | 3125 | A | A2M |
| Chain 1 | 56 | 3172 | C | OMC |
| Chain 1 | 57 | 3200 | G | OMG |
| Chain 1 | 58 | 3203 | A | 2MA |
| Chain 1 | 59 | 3213 | U | OMU |
| Chain 1 | 60 | 3223 | G | OMG |
| Chain 1 | 61 | 3227 | A | A2M |
| Chain 1 | 62 | 3301 | A | A2M |

### SSU modifications

| Chain | Sl. no | Nucleotide no. | Nucleotide | Modification type |
|---|---|---|---|---|
| Chain a | 1 | 91 | G | OMG |
| Chain a | 2 | 105 | A | A2M |
| Chain a | 3 | 161 | A | A2M |
| Chain a | 4 | 431 | A | A2M |
| Chain a | 5 | 735 | U | OMU |
| Chain a | 6 | 793 | A | A2M |
| Chain a | 7 | 944 | U | OMU |
| Chain a | 8 | 954 | A | A2M |
| Chain a | 9 | 1010 | A | A2M |
| Chain a | 10 | 1016 | A | A2M |
| Chain a | 11 | 1151 | G | OMG |
| Chain a | 12 | 1584 | U | OMU |
| Chain a | 13 | 1762 | C | OMC |
| Chain a | 14 | 1805 | C | 4OC |
| Chain a | 15 | 1807 | C | 5MC |
| Chain a | 16 | 1895 | G | OMG |
| Chain a | 17 | 1920 | C | 4AC |
| Chain a | 18 | 1928 | A | MA6 |
| Chain a | 19 | 1929 | A | MA6 |
